# The RNA-binding protein, Rasputin/G3BP, enhances the stability and translation of its target mRNAs

**DOI:** 10.1101/2020.01.21.913079

**Authors:** John D. Laver, Jimmy Ly, Allison K. Winn, Angelo Karaiskakis, Sichun Lin, Kun Nie, Giulia Benic, Nima Jaberi-Lashkari, Wen Xi Cao, Alireza Khademi, J. Timothy Westwood, Sachdev S. Sidhu, Quaid Morris, Stephane Angers, Craig A. Smibert, Howard D. Lipshitz

## Abstract

G3BP RNA-binding proteins are important components of stress granules (SGs). Here we analyze the role of *Drosophila* G3BP, Rasputin (RIN), in unstressed cells, where RIN is not SG associated. Immunoprecipitation followed by microarray analysis identified over 550 mRNAs that copurify with RIN. The mRNAs found in SGs are long and translationally silent. In contrast, we find that RIN-bound mRNAs, which encode core components of the transcription, splicing and translation machinery, are short, stable and highly translated. We show that RIN is associated with polysomes and provide evidence for a direct role for RIN and its human homologs in stabilizing and upregulating the translation of their target mRNAs. We propose that when cells are stressed the resulting incorporation of RIN/G3BPs into SGs sequesters them away from their short target mRNAs. This would downregulate the expression of these transcripts, even though they are not incorporated into stress granules.

## INTRODUCTION

Post-transcriptional regulation (PTR) plays a key role in control of gene expression in all cell types. In the cytoplasm of a cell, PTR is exerted at the level of mRNA stability, translation and subcellular localization (Bovaird et al., 2018; Tutucci et al., 2018). PTR is achieved by RNA-binding proteins (RBPs) and small RNAs such as microRNAs (miRs), which act as specificity factors that modulate the interaction of mRNAs with the cellular machinery that localizes, translates and degrades mRNAs (Achsel and Bagni, 2016; Iadevaia and Gerber, 2015; Van Treeck and Parker, 2018).

PTR is particularly important in early animal embryos, where maternally provided mRNAs and proteins control developmental events prior to transcriptional activation of the embryo’s genome (Tadros and Lipshitz, 2009; Vastenhouw et al., 2019). In several model animals, including *Drosophila*, the transfer of control from maternal products to those synthesized by the embryo’s own genome––the maternal-to-zygotic transition (MZT)–– is very rapid, occurring over a matter of hours, thus facilitating studies of the mechanisms and functions of PTR. For example, it has been shown that the *Drosophila* RBP, Smaug (SMG), which binds to specific stem-loop structures in its target mRNAs to repress their translation and trigger their degradation (Aviv et al., 2006; Aviv et al., 2003; Chen et al., 2014; Semotok et al., 2005; Semotok et al., 2008; Smibert et al., 1999; Smibert et al., 1996), is essential for repression/clearance of hundreds of maternal mRNAs and for timely activation of the zygotic genome (Benoit et al., 2009; Laver et al., 2015b; Luo et al., 2016; Tadros et al., 2007). SMG is not the only negative regulator of maternal transcripts in *Drosophila*; additional RBPs (e.g., Brain tumor, Laver et al., 2015a) or miRs (e.g., miR-309, Bushati et al., 2008) function in maternal mRNA clearance.

PTR also serves as a rapid response to cellular stress. Under stress conditions, cells shut down translation of many mRNAs while up-regulating transcription and/or translation of sets of protein chaperones that maintain basal cellular integrity. Repression occurs, at least in part, in membraneless organelles known as stress granules (SGs) (Harvey et al., 2017; Panas et al., 2016; Protter and Parker, 2016; Van Treeck and Parker, 2018). SGs are thought to contain transcripts that are stalled in translation initiation (Anderson and Kedersha, 2009a, b; Buchan and Parker, 2009; Kedersha and Anderson, 2009) and recent global analyses have shown that SGs are enriched for long transcripts (Khong et al., 2017; Namkoong et al., 2018).

An important component of SGs is the RBP, G3BP, which is conserved throughout eukaryotes. Mammals have two genes, G3BP1 and G3BP2, whereas in *Drosophila* there is a single gene, *Rasputin* (*rin*). In human cells, G3BPs are necessary for SG formation and, if over-expressed, they are sufficient to induce SGs even in the absence of stress (Tourriere et al., 2003). RIN is necessary for SG formation in the *Drosophila* S2 tissue culture cell line and, while RIN or G3BP overexpression can induce SGs in human cells, this is not the case in S2 cells (Aguilera-Gomez et al., 2017). RIN and G3BPs interact with several of the same protein partners under both stress and non-stress conditions; these include Caprin (CAPR), FMRP (FMR1 in *Drosophila*) and UPA2 (Lingerer [LIG] in *Drosophila*) (Aguilera-Gomez et al., 2017; Baumgartner et al., 2013; Costa et al., 2013; Jain et al., 2016; Kedersha et al., 2016; Markmiller et al., 2018; Youn et al., 2018). However, the roles of these interactors in SG assembly or disassembly can vary under different conditions. For example, in human cells, G3BP interaction with CAPR nucleates SG formation under oxidative stress (Solomon et al., 2007; Tourriere et al., 2003) but G3BP and CAPR are not required for SG formation under osmotic stress (Kedersha et al., 2016).

The roles of RIN/G3BPs in unstressed cells have received less attention than their roles upon stress. Multiple functions have been attributed to G3BPs (reviewed in Alam and Kennedy, 2019), including transcript destabilization/repression (e.g. *c-myc*, *BART*, *CTNNB1*, *PMP22*, *HIV-1*, *miR-1*), transcript stabilization (e.g., *Tau*, *SART3*), subcellular transcript localization (e.g., *Twist1*), and transcript sequestration into virus-induced foci (e.g., *HIV-1*). In *Drosophila*, mutations in the *rin* gene cause severe defects in oogenesis, mutant females lay few eggs and those that are laid fail to hatch (Costa et al., 2013). *rin* mutations can also result in tissue patterning and growth defects (Baumgartner et al., 2013; Pazman et al., 2000). Despite RIN’s essential function in the fly life cycle, there have been no analyses of the RIN-bound transcriptome or RIN’s global role in gene regulation, nor are the molecular mechanisms that underlie RIN function known.

To better understand the function of RIN/G3BP in unstressed cells, we have carried out global analyses of the RIN-associated proteome and transcriptome in early *Drosophila* embryos. Using an anti-RIN synthetic antibody that we isolated from a phage-displayed library of fragments antigen binding (Fab) (Na et al., 2016; Persson et al., 2013), we immunoprecipitated (IPed) RIN and then carried out mass spectrometry (IP-MS) to confirm IP of RIN and to identify RIN’s partner proteins. Interactions were found with several RBPs previously shown to interact with G3BP/RIN (e.g., Caprin, FMR1, and Lingerer), consistent with IP of a biologically relevant RIN-containing complex or complexes. RNA-dependent interactions with RIN were found for both small and large ribosomal subunit proteins, suggesting that RIN may be polysome associated, a fact we confirmed using polysome gradients. By coimmunoprecipitating RIN together with bound mRNAs followed by microarray analysis (RIP-chip) we identified hundreds of *in vivo* target transcripts in embryos, which are characterized by two features: they are short and are enriched for a RIN-binding motif that was previously identified *in vitro* (Cook et al., 2011; Ray et al., 2013). RIN-associated mRNAs are enriched for Gene Ontology (GO) terms for core components of the transcription, splicing and translation machinery as well as of mitochondria. RIN’s endogenous targets in early embryos are more stable and more highly translated than co-expressed unbound transcripts. Their shorter length and higher rates of translation contrast with the behaviour of mRNAs associated with SGs. Consistent with a role for RIN as a positive regulator of transcript stability, in *rin* mutants the abundance of several highly bound target mRNAs is reduced relative to controls. Using a heterologous RNA-binding domain to tether RIN, G3BP1 or G3BP2 to a luciferase reporter mRNA in S2 tissue culture cells, we confirmed that RIN/G3BP increases the stability and/or translation of bound transcripts in the absence of stress.

Our data support a conserved function for G3BP proteins as potentiators of the translation and stability of their target transcripts. We speculate that stress-dependent recruitment of G3BPs/RIN into SGs may serve as a mechanism to down-regulate gene expression both directly, by removing RIN from its endogenous target mRNAs, as well as indirectly, through reduced transcription, splicing and translation.

## RESULTS

### Expression of RIN in early embryos

We analyzed endogenous RIN expression in early embryos by western blots and whole-mount immunofluorescence using a previously published rabbit polyclonal antibody (Aguilera-Gomez et al., 2017). The expression level of RIN does not change significantly through the time course examined (0–4 hr; Figure S1A, B). RIN is enriched in the cortex of the early embryo (Figures S1C, D and S2) and is concentrated apical to the nuclei at the syncytial blastoderm stage (Figures S1E and S2), and is not present in foci. We compared the distribution of RIN with a known interacting protein, FMR1 (Monzo et al., 2006), and found that their distributions overlap in the cortex (Figure S1C-E).This pattern for RIN is consistent with previously published data on the *Drosophila* ovary, showing that RIN is enriched in the cortical cytoplasm of the nurse cells and the oocyte (Costa et al., 2013).

### RBPs, translation factors and ribosomal proteins co-purify with RIN, which exhibits polysome association

Our next goal was to identify protein partners and mRNA targets of RIN in early embryos. An anti-RIN synthetic antibody, D072, was generated against its NTF2-like domain (see Figure 1A and Methods for details) and was used to IP endogenous RIN from 0–3 h old embryos followed by mass spectrometry (LC-MS/MS). To differentiate proteins that associate with RIN via protein-protein interactions versus through co-binding to RIN’s target RNAs, IPs were performed either in the presence (+) or absence (–) of RNase A. In both sets of IPs, RIN was among the most abundant proteins identified and was greatly enriched in anti-RIN IPs compared to those performed with a control synthetic antibody, C1 (Laver et al., 2012; Laver et al., 2013; Laver et al., 2015a; Na et al., 2016) (Figure 1B).

**Figure 1.**
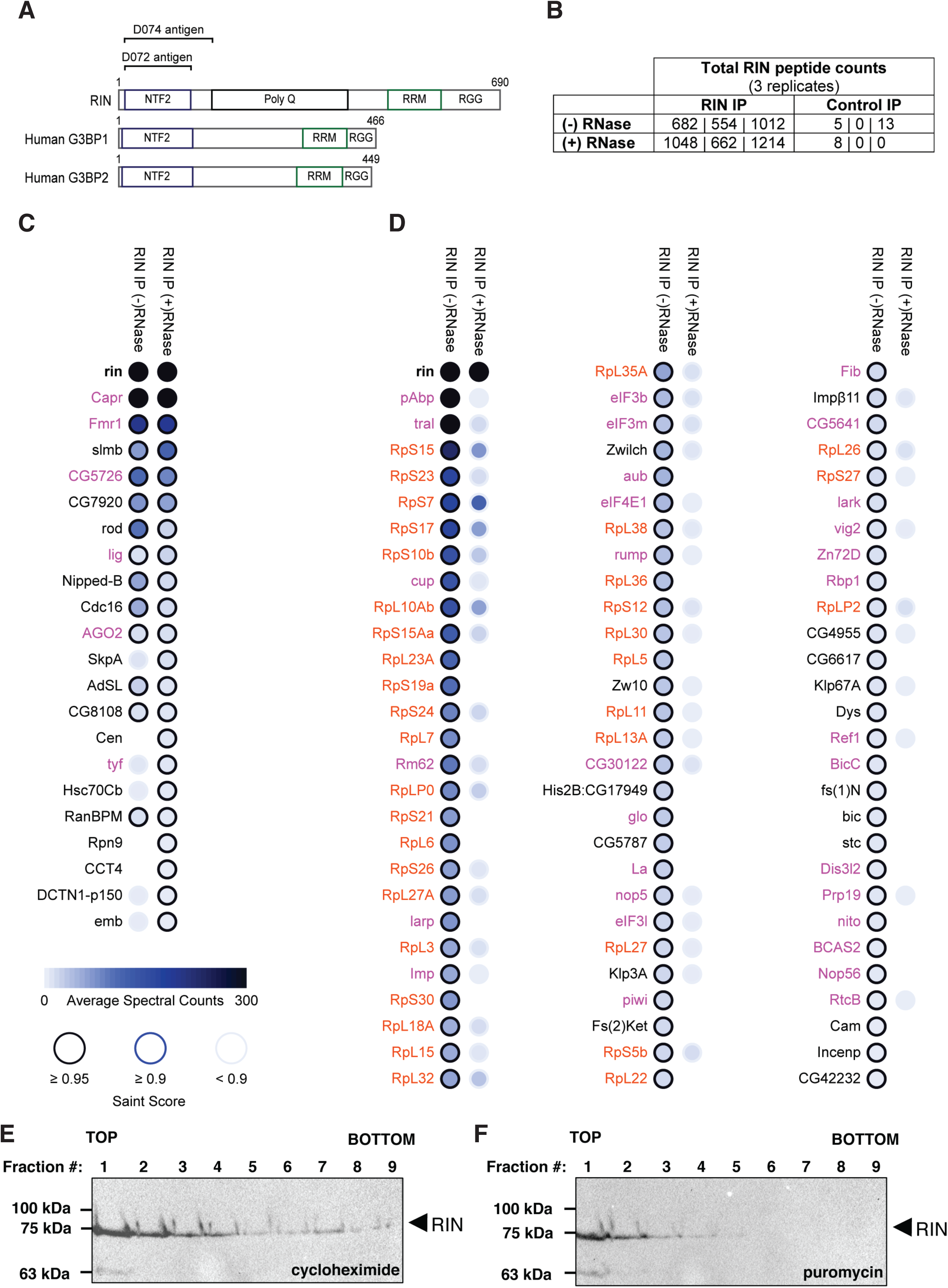
RIN interacts with multiple RNA-associated proteins in early embryos and is polysome associated. **(A)** The domain structure of *Drosophila* RIN and its human orthologs G3BP1 and G3BP2. Synthetic antibodies used in this study were generated against antigens consisting of regions of RIN encompassing the NTF2 domain, as indicated. **(B)** Total RIN peptide counts in IPs using the anti-RIN antibody D072, or the C1 control antibody, demonstrating that RIN is efficiently and specifically immunoprecipitated from early embryos. (**C** and **D**) RIN IP-MS results depicted as dotplots generated using the ProHits-viz web server (Knight et al., 2017). Shown are proteins identified as RIN interactors by analysis of IP-MS data using SAINT, defined as those with a SAINT score ≥ 0.95 and BFDR ≤ 0.01. In **(C)** are proteins identified as RNA-independent interactors (scored as significant in IPs (+) RNase), and in **(D)** are proteins identified as RNA-dependent interactors (scored as significant in IPs (-) RNase but not (+) RNase). The shade of the dot fill represents protein abundance in the RIN IPs minus control IPs, based on total peptide counts, and the shade of the dot outline represents the SAINT score, as indicated in the legend at the bottom of panel **(C)**. Protein names highlighted in magenta indicate those that are known RBPs, mRNP complex components, or translation factors, and orange highlighting indicates ribosomal proteins. **(E, F)** Embryo extract treated with either cycloheximide **(E)** or puromycin **(F)** was fractionated on sucrose gradients, with the resulting fractions assayed for RIN via western blot.

We performed Significance Analysis of Interactome (SAINT, Choi et al., 2011), and defined RIN-interactors as proteins that were significantly enriched in the RIN IPs with a SAINT score ≥ 0.95 and a Bayesian false discovery rate (BFDR) ≤ 0.01. In addition to RIN itself, 21 proteins were enriched in the (+) RNase IPs and 96 proteins were enriched in the (–) RNase IPs (Figure 1C and D; Table S1). Of the 21 RNA-independent interactors, 13 were also positives in the (–) RNase IPs while eight were not. However, based on total peptide counts, five out of these eight were enriched in one of three (–) RNase IP biological replicates; the stronger enrichment in the (+) RNase samples may have been due to decreased background in the control IPs in the presence of RNase.

Consistent with a role for RIN in post-transcriptional regulation in the early embryo, of the total of 104 proteins identified in our RIN IPs, 68% (71) are known to be involved in post-transcriptional processes (as annotated by FlyBase Release FB2019_06; http://flybase.org/): 36 of them are known RBPs, mRNP complex components or translation factors (Figure 1C and D, highlighted in magenta), while 35 are ribosomal proteins (Figure 1C and D, highlighted in orange), including 36% of the small subunit and 39% of the large subunit proteins (respectively, 14 of 39 and 21 of 54) (Marygold et al., 2007). Overall, 92% (66) of the RBPs and ribosomal proteins that co-purified with RIN did so in an RNA-dependent manner, including all of the 40S and 60S subunit proteins.

Previous studies have identified sets of RIN/G3BP-interacting proteins in other contexts and species, and we therefore compared these to our IP-MS data to ask whether any of these interactions are conserved in *Drosophila* embryos. In *Drosophila* ovaries, RIN has been shown to be in an RNase-resistant complex with ORB (Costa et al., 2005); ORB is absent from embryos (Hafer et al., 2011) and therefore did not co-IP in our experiments. Another mass-spectrometric analysis, which used Dorsal (DL) as a negative control, identified additional RIN-interacting proteins in ovaries (Costa et al., 2013). In that study, of the proteins that they found to IP with RIN but not DL, eight were also present on our early embryo lists: three were RNA-independent (FMR1, Lingerer [LIG], Twenty-four [TYF]) and four were RNA-dependent (PABP, RpS24, RpL11, RpL13A). An additional nine proteins on our early embryo lists were found in both their RIN and their DL IPs: one (CAPR) was on our RNA-independent list, and eight were on our RNA-dependent list (Trailer-hitch [TRAL], Cup, Fibrillarin [FIB], RpL3, RpL15, RpLP0, RpLP2, RpS5).

Human G3BPs interact with many of the same proteins in both unstressed and stressed cells (Jain et al., 2016; Markmiller et al., 2018; Youn et al., 2018). We, therefore, compared our list of RIN interactors with G3BP-interactors as well as with SG components identified in recent global proteomic analyses in human cells (Jain et al., 2016; Markmiller et al., 2018; Solomon et al., 2007; Youn et al., 2018) and budding yeast (Jain et al., 2016) (Table S2). 24% (5/21) of RIN’s RNA-independent and 25% (24/96) of the RNA-dependent interactors had been identified as G3BP interactors in the human datasets. Regarding RIN’s RNA-independent interactors, all of the human datasets included CAPR, FMR1 and LIG and one study (Jain et al., 2016) found two additional proteins that were on our RNA-independent list (HSC70Cb, DCTN-p150). None of our RNA-independent interactors overlapped with budding yeast SG components while seven of the proteins on our RNA-dependent list overlapped (PABP, TRAL, LARP, RUMP, eIF3b, eIF4E1, RpS30). Finally, regarding our RNA-dependent list, 84% of the proteins that overlapped with the human or yeast datasets are RBPs/translation factors (21/25).

G3BP interacts with the 40S (but not the 60S) ribosomal subunit in the context of SGs and their formation (Kedersha et al., 2016). Interestingly, our data showed that RIN interacts with multiple proteins from both the 40S and 60S subunits in early embryos, which lack SGs. Since these RIN interactions were RNase sensitive, one possibility is that, in early embryos, RIN binds to mRNAs that are actively translated, and thus the ribosomes transiting these mRNAs co-purify with RIN. To assess whether RIN is associated with polysomes, we ran early embryo extracts on sucrose gradients treated with cycloheximide (which stabilizes the interaction of ribosomes with mRNAs) or puromycin (which disrupts mRNA-ribosome associations), followed by western blotting with an anti-RIN antibody (Aguilera-Gomez et al., 2017) (Figures 1E and S3). These experiments showed that a subset of RIN is polysome-associated.

Taken together, our data confirm that numerous RIN/G3BP-partner protein interactions occur under both non-stress and stress conditions, and are conserved in *Drosophila*, yeast and human cells. Furthermore, unlike in stressed cells, in early embryos RIN is found on polysomes. A possible role for RIN in mRNA translation and stability is investigated below.

### RIN associates with hundreds of mRNA species in the early embryo

To identify mRNAs associated with RIN in 0–3 h embryos, we used the D072 anti-RIN synthetic antibody to carry out RNA co-immunoprecipitations followed by microarray analysis (RIP-Chip). Significance Analysis of Microarrays (SAM) (Tusher et al., 2001) identified the mRNAs encoded by 566 genes that were enriched at least two-fold and with an FDR < 5% in RIN IPs compared to control IPs performed with the C1 antibody. We defined these as RIN-associated mRNAs (Figure S4; Table S3).

Given that no global analyses of endogenous mRNAs associated with *Drosophila* RIN had been carried out previously, we validated our set of RIN-associated transcripts in two ways. First, we performed two additional biological replicate RIPs using the anti-RIN antibody (D072) and control antibody described above, after which reverse transcription followed by quantitative polymerase chain reaction (RT-qPCR) was used to quantify five highly enriched RIN-associated mRNAs and four mRNAs not associated with RIN according to our RIP-Chip experiments. The five putative RIN-associated mRNAs were two-to seven-fold more enriched in the RIN versus control RIP-RT-qPCR than were the non-target mRNAs (Figure S5). Second, we generated an additional synthetic antibody, designated D074, against the NTF2-like domain of RIN (Figure 1A), which we used to perform RIP-Chip, and compared the average fold-enrichment of transcripts in the initial anti-RIN D072 RIP-Chip to the fold-enrichments in the RIP-Chip performed with D074. The results were consistent with the initial RIP-Chip data, with Spearman correlation rho value for the comparison of fold enrichments of 0.79 (*P* < 10^-15^) (Figure S6).

### RIN-associated mRNAs are short and enriched for the RIN RRM’s *in vitro* binding-motif

Next we assessed the properties of RIN-associated transcripts. Strikingly, we found that RIN-associated mRNAs in early embryos are significantly shorter than co-expressed unbound transcripts with the Area Under the Receiver Operating characteristic (AUROC) = 0.83 (*P*<10^-10^; Figure 2A). The ROC graphs the diagnostic ability of a binary classifier by plotting the true-positive rate versus the false-positive rate; no diagnostic ability gives an AUROC of 0.5 while perfect ability gives an AUROC of 1.0. For the Mann-Whitney *U* test, the AUROC is equivalent to the *U* statistic divided by the product of the numbers of targets and non-targets.

**Figure 2.**
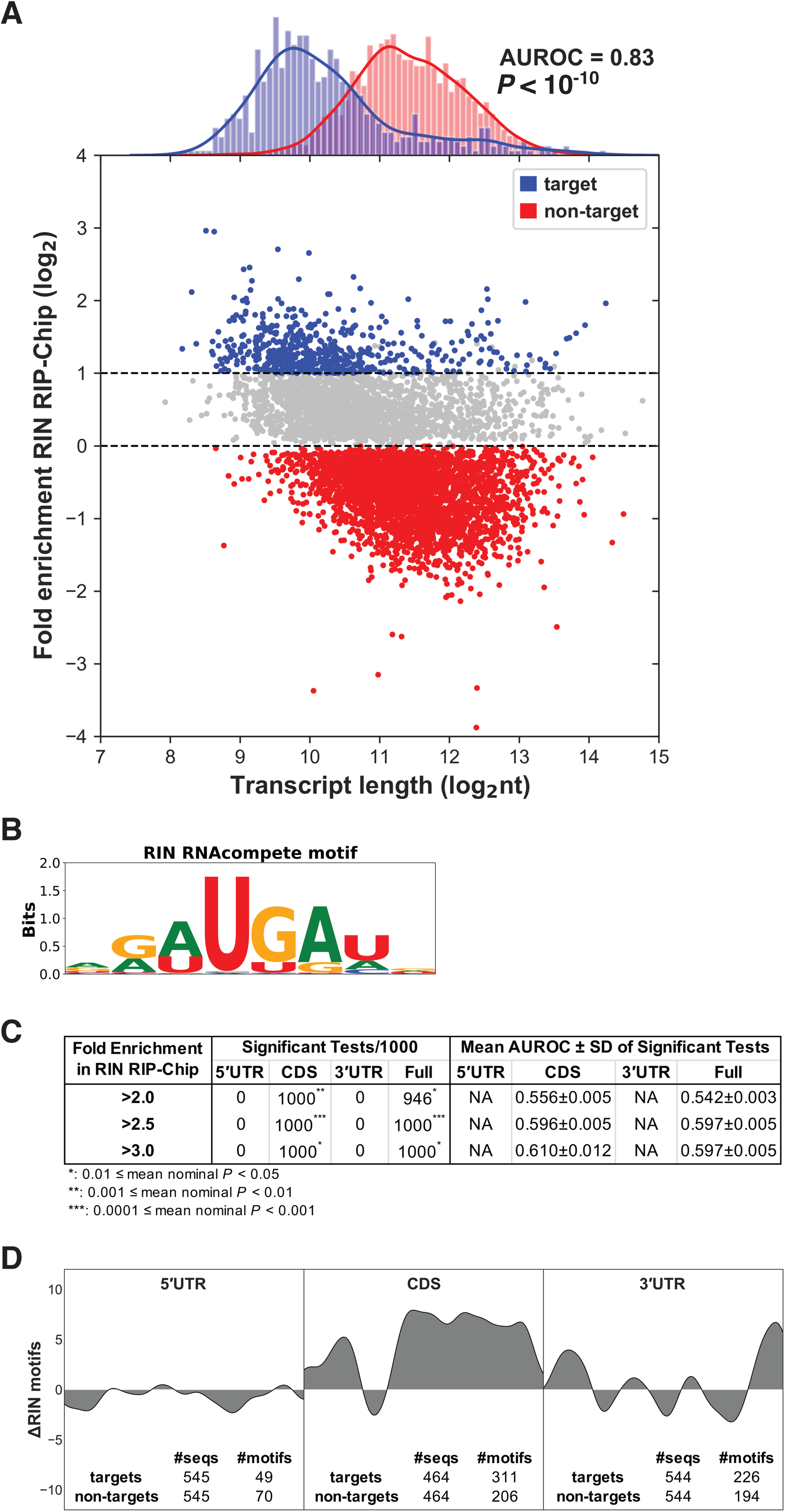
RIN-associated transcripts are short and enriched for the RIN RRM’s *in vitro* binding-motif. **(A)** Plot showing the relationship between transcript length and association with RIN, for all genes with transcripts represented on the microarray that were defined as expressed in early embryos. Transcript length for each gene is taken to be the length of the longest annotated transcript isoform, and RIN association is represented by the fold-enrichment in the RIN RIP-Chip versus control RIP-Chip of the most highly enriched probe set on the microarray for each gene. Genes encoding RIN-associated transcripts (defined as enriched > 2-fold in RIN RIP versus C1 RIP with FDR < 5%) are highlighted in blue. Co-expressed RIN non-targets (defined as log_2_ fold-enrichment in RIN versus C1 RIPs < 0) are highlighted in red. Gene length distributions of RIN targets and non-targets are shown in the density plots at the top. AUROC and *P* value of Mann-Whitney U test on transcript lengths of RIN targets and non-targets is shown on the plot. **(B-D)** The *in vitro* RIN motif from RNAcompete (Ray et al., 2013) **(B)** is significantly enriched in the coding sequences of RIN-associated mRNAs **(C, D)**. **(C)** Table showing the number, AUROC and range of nominal *P* values of significant tests out of 1000 trials with randomly selected targets and length-paired non-targets. **(D)** Plot showing the difference in number of RIN motifs in the 3’UTR, CDS and 5’UTR of RIN-associated transcripts versus randomly selected co-expressed length-paired RIN non-targets for each region. The total number of RIN motifs (defined as sites with “hit score” > 0.001) in targets and non-targets for each region are grouped into 20 bins. The difference between the two Gaussian kernels fitting the motif distribution of targets and non-targets for each region is displayed on the plot. The locations of motifs are represented by the relative location of the first nucleotide in the corresponding region.

To assess whether enrichment of short mRNAs might be an artefact of our RIP-chip method, we reanalyzed our previously published data for three other RBPs – Brain tumor, Pumilio, and Staufen (endogenous Staufen and transgenic GFP-Staufen) – for which we used the same method and microarray platform (Laver et al., 2013; Laver et al., 2015a). For all four RIP-Chip datasets (Table S4) – in contrast to the RIN RIP-Chip dataset – there was either enrichment for long mRNAs relative to co-expressed unbound transcripts (Figure S7A, B, D) or no length difference (Figure S7C). We conclude that short mRNAs do indeed co-purify with RIN.

To identify features other than transcript length that distinguish RIN-associated mRNAs, we performed *de novo* motif discovery using #ATS, a discriminative tool that searches for sequence motifs that are enriched in targets versus non-targets, in computationally predicted single-stranded regions (Li et al., 2010). We failed to discover a consistent and significant motif in RIN targets compared to length-matched co-expressed non-targets (Table S5).

RIN and G3BP-binding motifs have been reported from *in vivo* (Edupuganti et al., 2017) and *in vitro* (Cook et al., 2011; Ray et al., 2013) studies. G3BP-family proteins are characterized by a single RNA-recognition motif (RRM), which was used for the *in vitro* analyses (Cook et al., 2011; Ray et al., 2013). To assess whether any of these motifs were enriched in RIN-associated mRNAs, we used position frequency matrices (PFMs) from those studies together with RNAplfold, where the latter was used to define motif matches that were single-stranded and thus available for RBP binding (Bernhart et al., 2006; Li et al., 2010). For each motif, we tested its ability to distinguish RIN-associated (positive) transcripts from length-paired, co-expressed, non-RIN-associated (negative) transcripts, by ranking transcripts with the maximum “hit score” achieved for that motif, across all subsequences of a transcript (i.e., the best possible subsequence resembling the motif sequence in a context that is predicted to be single-stranded). We performed this test using three RIN fold-enrichment (FE) cut-offs (>2, >2.5 and >3) and four transcript regions (5’UTR, CDS, 3’UTR, full mRNA), and repeated each FE and region pair 1000 times with length-matched negatives randomly selected each time. We assessed how well the four motifs performed on samples from each FE cut-off and transcript region by the number of Mann-Whitney *U* tests that were significant (with *P* < 0.05), and the mean and standard deviation of the AUROC. We found that the *in vitro* RIN motif (Figure 2B) was significantly enriched in RIN-associated full mRNA sequences with the mean AUROC increasing with FE from 0.54 at FE > 2.0 to 0.61 at FE > 3.0 (Figure 2C, Table S6). The motif was most enriched in the open reading frame, with the mean AUROC increasing from 0.56 at FE > 2.0 to 0.61 at FE > 3.0 (Figure 2C), and 105 more motifs found in targets versus length-matched non-targets (Figure 2D). None of the motifs for the human G3BPs––defined either *in vitro* or *in vivo*––was enriched (Table S6).

We conclude that the major feature of RIN-binding *in vivo* is short transcript length (AUROC = 0.83) with an additional, more minor, contribution from the motif (AUROC 0.55 to 0.61).

### RIN-associated mRNAs are stable, translated, and depleted for Smaug recognition elements

To gain insight into the effect RIN might have on the expression of its target mRNAs, we examined datasets that define the translational status (determined by ribosome occupancy) and abundance of the *Drosophila* transcriptome during late oogenesis and early embryogenesis, and asked whether RIN-association correlated with any particular patterns of post-transcriptional outcome. We focused on the data from mature (Stage 14) oocytes and two embryonic stages that overlap with the time window in which we carried out our RIP-chip: 0–1 h and 2–3 h post-fertilization First, we examined the abundance of RIN-bound mRNAs and of co-expressed, unbound mRNAs (Eichhorn et al., 2016). Whereas the median abundance of these two classes of transcripts was similar in mature oocytes and 0–1 h embryos, there was a striking change moving from 0–1 h embryos to 2–3 h embryos: non-target mRNAs significantly decreased in abundance (2.2-fold decrease in median RPKM, Wilcoxon rank-sum *P* < 10^-150^) whereas levels of RIN-associated transcripts increased significantly (1.4-fold increase in median RPKM, Wilcoxon rank-sum *P* < 10^-3^) (Figure 3A). This difference was maintained in 3–4 and 4–5 h embryos (Figure S8). Since RIN-associated mRNAs are short, we also carried out abundance analyses relative to length-matched non-targets and found that RIN-bound mRNAs are more stable at 2–3 hours even when length is taken into account (Figures 3B, S9).

**Figure 3.**
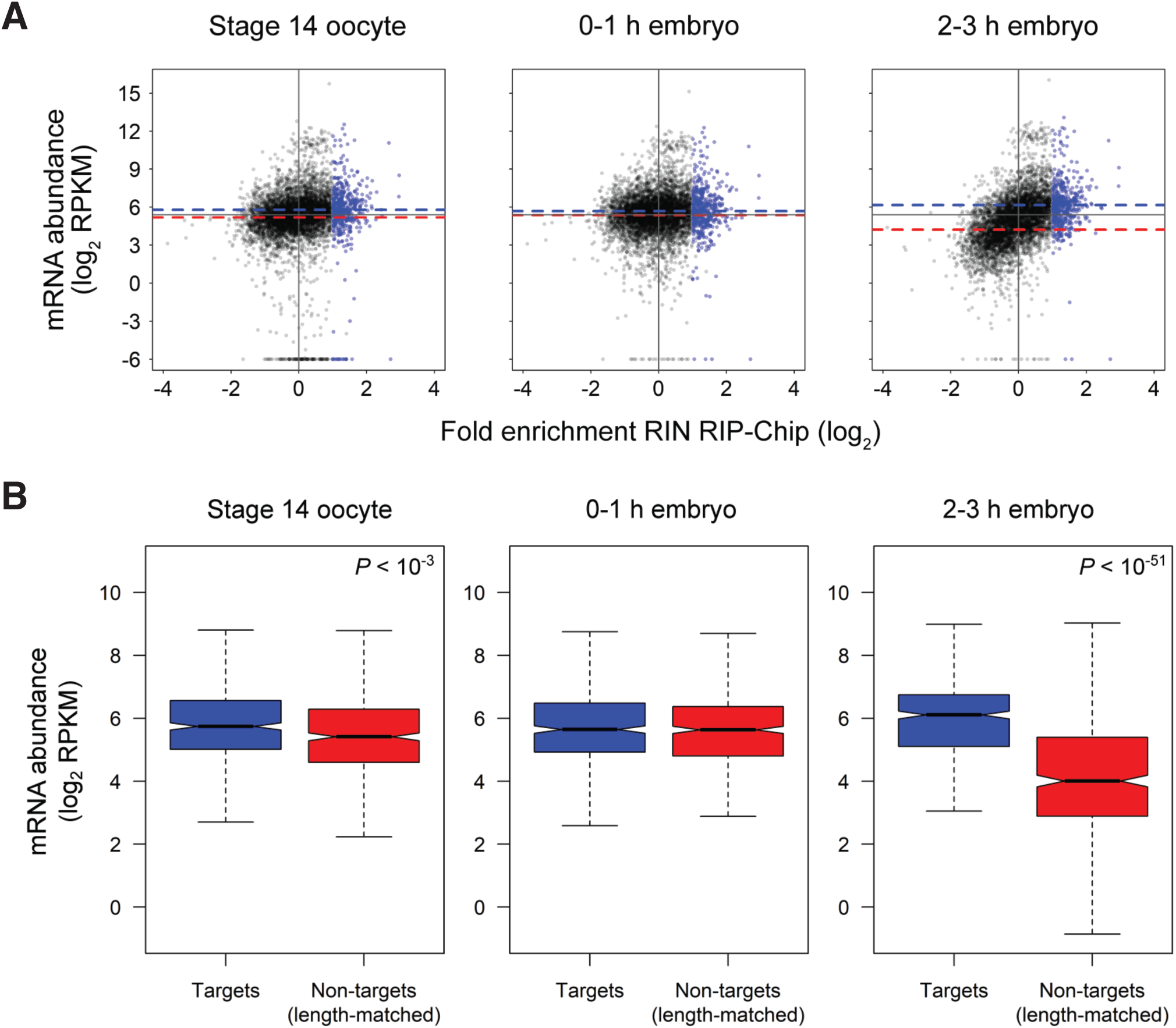
RIN-associated transcripts are stable during the maternal-to-zygotic transition. **(A)** Plots showing the relationship between fold-enrichment in the RIN RIP-Chip and mRNA levels, measured by Eichhorn et al. (2016), during timepoints spanning the maternal-to-zygotic transition. RIN-associated transcripts as defined in this manuscript are highlighted in blue. Dashed blue horizontal lines indicate median value of mRNA levels for RIN-associated transcripts at each timepoint, and dashed red horizontal lines indicate median values for RIN non-targets (defined as log_2_ fold-enrichment in RIN versus C1 RIPs < 0) at each timepoint. For all timepoints, solid dark-grey horizontal lines indicate the median values of mRNA levels in stage 14 oocytes, as a reference, and solid vertical lines indicate no enrichment in the RIN RIP-Chip. Points are shown for all genes measured in both this study and Eichhorn et al., with the RIP-Chip fold-enrichment value for each gene represented by the most highly enriched probe set on the microarray for that gene. **(B)** Boxplots comparing mRNA levels, measured by Eichhorn et al. (2016) as in (A), of RIN-associated transcripts (“Targets”; blue) and a set of randomly selected, length-matched, co-expressed non-RIN-targets (red), for each time point depicted in (A). For comparisons with a statistically significant difference in mRNA levels between targets and length-matched non-targets, Wilcoxon rank-sum *P* values are indicated in the top right of the plots.

The first three hours of embryogenesis encompass most of the *Drosophila* maternal-to-zygotic transition, during which many maternally supplied mRNAs are degraded and the zygotic genome becomes transcriptionally active (Tadros and Lipshitz, 2009; Vastenhouw et al., 2019). Transcripts present in the early embryo have been classified according to their maternal and/or zygotic origin, as well as their patterns of stability or decay (De Renzis et al., 2007; Tadros et al., 2007; Thomsen et al., 2010). To understand how the abundance of RIN-associated transcripts described above reflects changes in levels of maternal or zygotic mRNAs, we compared RIN-associated mRNAs to these various classes. RIN-associated transcripts were enriched for two of these classes: maternally supplied transcripts that remain stable through the MZT and are not transcribed in the embryo (Class I in Thomsen et al., 2010); Fisher’s exact test *P* < 10^-11^, odds ratio 2.0), and maternally supplied transcripts that remain stable and are also zygotically transcribed (‘Stable + Transcription’ in Thomsen et al., 2010) (Fisher’s exact test *P* < 10^-27^, odds ratio 3.2). However, RIN-associated mRNAs were strongly depleted for maternally supplied transcripts that are degraded (Classes II-V in Thomsen et al., 2010) (Fisher’s exact test *P* < 10^-48^, odds ratio 0.24) including those that are degraded and subsequently re-transcribed (Class III in Thomsen et al., 2010) (Fisher’s exact test *P* < 10^-7^, odds ratio 0.20). Consistent with these results, examination of two additional datasets, which defined mRNAs that are maternally supplied and degraded, revealed depletion of RIN-associated mRNAs among transcripts that are degraded in unfertilized eggs (Tadros et al., 2007) (Fisher’s exact test *P* < 10^-17^, odds ratio 0.26) or in early embryos (De Renzis et al., 2007) (Fisher’s exact test *P* < 10^-54^, odds ratio 0.12). These comparisons therefore indicate that the dynamics of RIN-associated transcript levels in early embryos, described above, largely reflect RIN binding to stable, maternally supplied transcripts.

Next, to assess the translational efficiency of RIN-associated mRNAs we used a dataset that measured this parameter using ribosome ‘footprinting’ (Eichhorn et al., 2016) This revealed that RIN-associated mRNAs had significantly higher translational efficiency than co-expressed, non-target mRNAs in both mature oocytes and early embryos, suggesting a possible role for RIN in potentiating translation. The largest and most significant difference was in 0–1 h embryos where the median translational efficiency of RIN-associated mRNAs was four-fold higher than that of non-target mRNAs (dashed blue and red lines, respectively, in Figure 4A; Wilcoxon rank-sum *P* < 10^-100^). Whereas the translational efficiency of RIN-associated mRNAs showed a significant increase moving from mature oocytes to 0–1 h embryos (1.6-fold increase, Wilcoxon rank-sum *P* < 10^-17^), the translational efficiency of co-expressed, non-target mRNAs decreased over the same stages. At earlier stages of oogenesis (Stages 11–13) as well as at later embryonic time points (3–4 and 4–5 h), RIN-bound transcripts also consistently exhibited higher translational efficiency than co-expressed unbound transcripts (Figure S8). The positive correlation between an mRNA’s association with RIN and its translational activity was supported by comparisons to an additional dataset that used ribosome footprinting to assess transcriptome-wide mRNA translation in early embryos (Dunn et al., 2013) (Figure S10). Since RIN-associated mRNAs are short, we also carried out analyses of translational efficiency relative to length-matched non-targets and found that RIN-bound mRNAs have higher translational efficiency at all timepoints even when length is taken into account (Figure 4B, S11).

**Figure 4.**
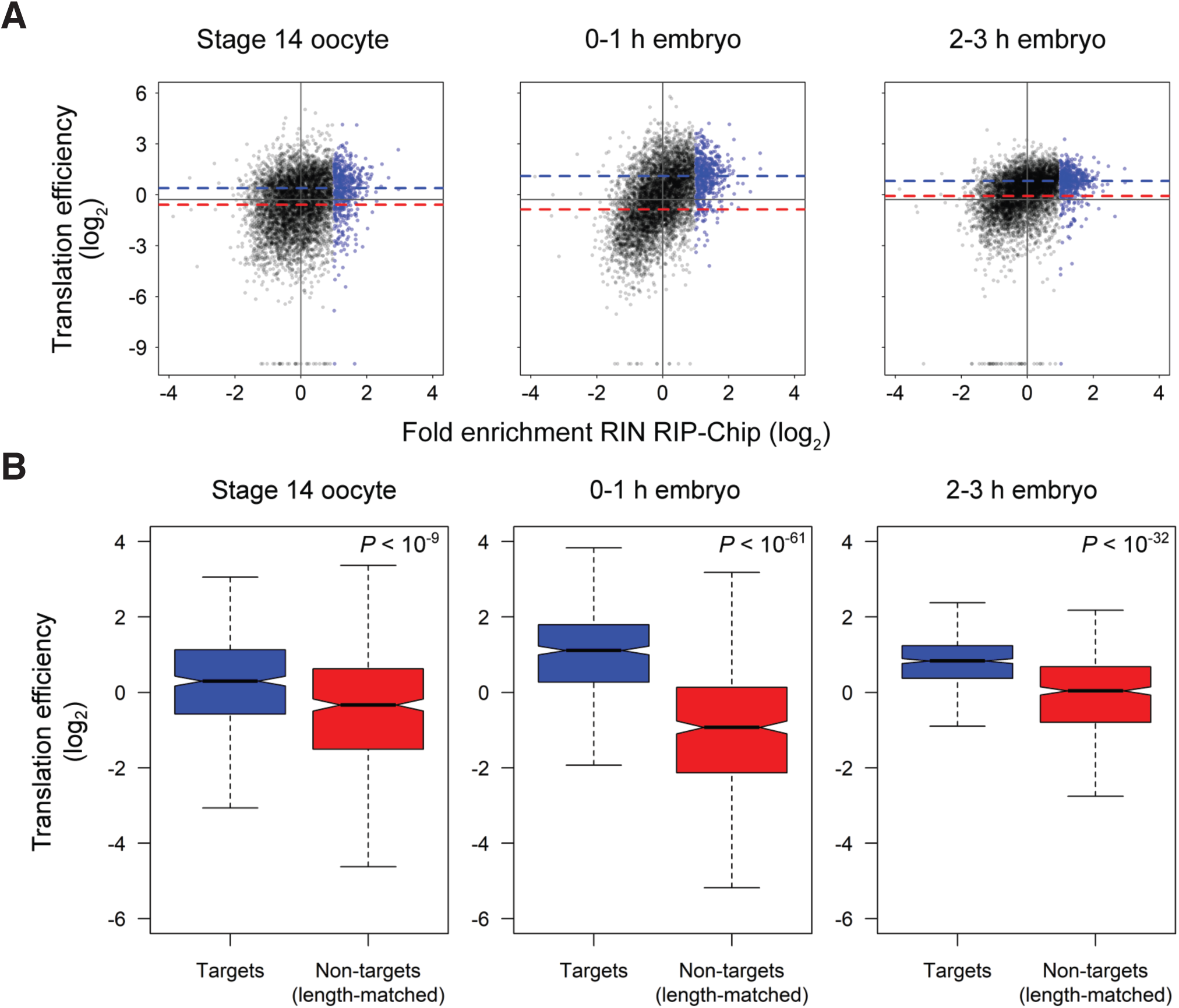
RIN-associated transcripts are highly translated during the maternal-to-zygotic transition. **(A)** Plots showing the relationship between fold-enrichment in the RIN RIP-Chip and translational efficiency, measured by Eichhorn et al. (2016), during timepoints spanning the maternal-to-zygotic transition. Color code is as in Figure 3 but with respect to translational efficiency rather than RNA abundance. For all timepoints, solid dark-grey horizontal lines indicate the median values of translational efficiency in stage 14 oocytes, as a reference, and solid vertical lines indicate no enrichment in the RIN RIP-Chip. Points are shown for all genes measured in both this study and Eichhorn et al., with the RIP-Chip fold-enrichment value for each gene represented by the most highly enriched probe set on the microarray for that gene. **(B)** Boxplots comparing translational efficiency, measured by Eichhorn et al. (2016) as in (A), of RIN-associated transcripts (“Targets”; blue) and a set of randomly selected, length-matched, co-expressed non-RIN-targets (red), for each time point depicted in (A). For comparisons with a statistically significant difference in translational efficiency between targets and length-matched non-targets, Wilcoxon rank-sum *P* values are indicated in the top right of the plots.

The Smaug RBP, which binds hairpin structures known as Smaug recognition elements (SREs), is a well-characterized negative regulator of both mRNA stability and translation (Chen et al., 2014; Pinder and Smibert, 2013; Semotok et al., 2005; Semotok et al., 2008; Smibert et al., 1996; Tadros et al., 2007). Since RIN-bound mRNAs are stable and translated in early embryos it was of interest to assess whether they might be depleted for SREs. We therefore calculated an SRE ‘score’ (Chen et al., 2014) for RIN-associated mRNAs relative to length-matched, co-expressed mRNAs, and found that RIN-bound mRNAs were strikingly depleted of SREs (Table S7).

A final feature that we examined was N^6^-methyladenosine (m^6^A) modification of RIN’s targets, since it has been reported that m^6^A “repels” G3BP1 binding (Edupuganti et al., 2017). If so, then RIN-bound transcripts would be expected to be depleted for m^6^A. Recently, m^6^A modifications have been mapped transcriptome-wide at several *Drosophila* embryonic stages (Kan et al., 2017). We, therefore, compared RIN-bound transcripts to this last study’s m^6^A transcriptome data from 0–45’, 45–90’ and 1.5–6 h embryos. In all cases and for all transcript regions (5’UTR, ORF, 3’UTR), there was neither significant enrichment nor significant depletion of m^6^A-modified transcripts among RIN’s targets (Fisher’s exact test *P* values ranged from 0.15 to 1). We conclude that there is no evidence for a role – either positive or negative – of m^6^A in determining RIN targets in early embryos.

Taken together, our data support a model in which RIN binds to maternally supplied transcripts, stabilizes them at the stage of the MZT when many other maternally supplied mRNAs are degraded and potentiates their translation.

### RIN-associated transcripts in the early embryo do not exhibit properties of stress granule transcriptomes

Human and yeast SG transcripts are long (Anderson and Kedersha, 2009a, b; Buchan and Parker, 2009; Kedersha and Anderson, 2009; Khong et al., 2017; Namkoong et al., 2018). As described above, in contrast, RIN-associated mRNAs in early embryos are significantly shorter than co-expressed unbound transcripts (Figure 2A). The high levels of translation of RIN-associated transcripts in the early *Drosophila* embryo also contrasts with the translationally silent transcripts found together with G3BPs in SGs (Anderson and Kedersha, 2009a, b; Buchan and Parker, 2009; Kedersha and Anderson, 2009).

To ask whether the RIN-associated transcripts are likely to be recruited to SGs under different conditions or, instead, represent a separate pool of RIN-associated mRNAs, we took advantage of recent reports that define SG transcriptomes in yeast and human cells. We compared our list of RIN-associated mRNAs to the *Drosophila* homologues of transcripts found to be either enriched in or depleted from human U2OS SGs (Khong et al., 2017): RIN-associated *Drosophila* mRNAs were significantly depleted for homologous SG-enriched transcripts (Fisher’s exact test *P* < 10^-7^, odds ratio 0.45) but enriched for SG-depleted transcripts (*P* < 10^-29^, odds ratio 3.2). Likewise, RIN-associated mRNAs were significantly depleted for transcripts homologous to those that undergo localization to RNA granules under ER stress (Fisher’s exact test *P* < 10^-4^, odds ratio 0.49), heat shock (Fisher’s exact test *P* < 10^-7^, odds ratio 0.55), or arsenite treatment (Fisher’s exact test *P* < 10^-8^, odds ratio 0.49) in NIH3T3 cells (Namkoong et al., 2018). The observed enrichment or depletion of these classes in RIN-bound transcripts is consistent with the fact that the fly homologs of SG-enriched or -depleted transcripts show the same length distribution as the human transcripts (Figure S12).

In addition to these comparisons, we compared the length and translational efficiency of the *Drosophila* homologs of mammalian mRNAs that are either enriched in or depleted from SGs. We found that, whereas there was no significant correlation of low translational efficiency and length of the fly homologs of SG-enriched transcripts, there was a significant negative correlation between length and translational efficiency of the fly homologs of SG-depleted transcripts (Figure S13).

Taken together, these analyses are consistent with the hypothesis that the transcripts with which RIN associates in early embryos are a separate pool from those typically found in SGs.

### RIN-associated mRNAs encode core gene expression and mitochondrial components

We next asked what biological and molecular functions RIN might control in early embryos, by searching for Gene Ontology (GO) terms enriched among the proteins encoded by the RIN-associated mRNAs, using the DAVID functional annotation tool (Huang da et al., 2009a; Huang da et al., 2009b) (Table 1; Table S8). This revealed striking enrichment for roles in regulation of gene expression and in mitochondrial function.

**Table 1.** Representative GO terms enriched among proteins encoded by RIN-associated mRNAs.

| Enriched function or complex | Representative GO terms | Fold-enrichment | FDR (%) |
| --- | --- | --- | --- |
| Oxidative phosphorylation | oxidative phosphorylation (GO:0006119) | 4.32 | $2.80 \times 10^{-5}$ |
| | respiratory chain complex (GO:0098803) | 4.21 | $2.92 \times 10^{-5}$ |
| Ribosome | mitochondrial ribosome (GO:0005761) | 3.43 | $3.63 \times 10^{-4}$ |
|  | cytosolic large ribosomal subunit (GO:0022625) | 2.65 | 5.81 |
| Ragulator complex | Ragulator complex (GO:0071986) | 11.02 | 0.44 |
| Transcription | DNA-templated transcription, initiation (GO:0006352) | 2.56 | 0.52 |
|  | histone acetylation (GO:0016573) | 2.35 | 9.11 |
| mRNA splicing | RNA splicing (GO:0008380) | 1.75 | 1.78 |

With regard to the regulation of gene expression, RIN’s targets included mRNAs encoding proteins involved in multiple levels of control, most notably transcription, pre-mRNA splicing, and translation. Among those involved in transcription were mRNAs encoding general transcription factors, including four subunits of the TFIID complex (*Taf1*, *Taf10b*, *Taf12*, *Taf13*), both of the subunits of the TFIIA complex (*TfIIA-S*, *TfIIA-L*), six subunits of the ‘mediator’ transcriptional co-activator complex (*MED4*, *7*, *11*, *21*, *26*, *28*), histone modifiers, and transcription factors with important roles in the activation of zygotic transcription, such as *Zelda* and *Stat92E*. mRNAs associated with RIN that encode splicing-related factors included a number of spliceosome subunits as well as regulators of pre-mRNA splicing, such as *transformer* and *transformer 2*. Finally, with regard to translation, RIN-associated mRNAs were enriched for those encoding protein components of both the cytosolic ribosome (12 large subunit, 2 small subunit) and the mitochondrial ribosome (15 large subunit, 8 small subunit). In addition, mRNAs that encode all five subunits of the ‘Ragulator’ complex were found to be RIN-associated. Ragulator regulates activation of the TOR pathway in response to amino acid availability (Bar-Peled et al., 2012; Sancak et al., 2010).

With regard to mitochondria, in addition to the transcripts encoding 23 mitochondrial ribosomal proteins, RIN-associated mRNAs were strongly enriched for transcripts encoding proteins involved in the mitochondrial electron transport chain. Indeed, RIN was associated with 30 nuclear-encoded mRNAs for proteins that are either components of the mitochondrial respiratory chain Complexes I to V or involved in their assembly. Additional mRNAs associated with RIN included those encoding components of the mitochondrial contact site and cristae-organizing system (MICOS) complex, which acts to maintain mitochondrial cristae and membrane architecture, and FIS1, which has a role in promoting mitochondrial fission.

We next examined each of the GO-term-enriched classes in detail. Strikingly, we found that the vast majority of transcripts in each category tended to be enriched in the RIN IP, albeit not to the level of statistical significance (i.e., tended to be to the right of ‘0’ on the x-axis; in Figure 5 compare transcripts associated with RIN, highlighted in blue, and transcripts that fell below the cut-off used to define RIN-associated mRNAs, highlighted in red). Notably, there was also a clear negative correlation between transcript length and RIN-binding in each GO-term class (i.e., the ‘blue’ transcripts tended to be shorter than the ‘red’ transcripts), indicating that the correlation of RIN-association with shorter transcript length is independent of the function of the bound mRNAs. Finally, the ‘red’ transcripts in each GO-term class tended to have similar translational efficiency and stability to the RIN-bound (‘blue’) transcripts in that class, albeit again not to as great an extent (Figure S14. These results suggest that RIN’s role in binding and regulating mRNAs that encode core components of transcription, splicing, translation and mitochondria is more pervasive than predicted only from examination of its most highly bound targets.

**Figure 5.**
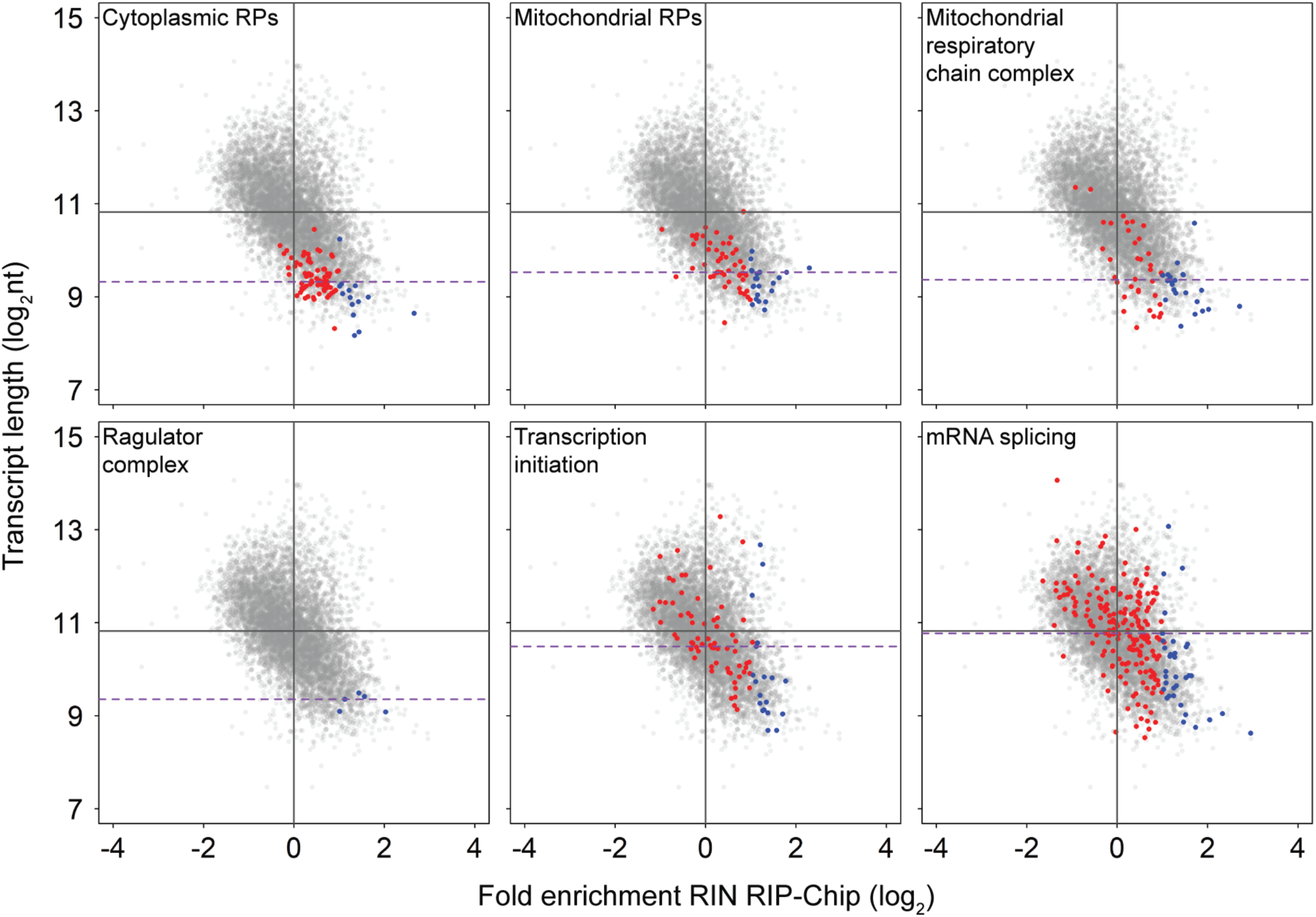
RIN association with mRNAs is correlated with shorter transcript length for all of the enriched GO-term categories. Plots showing, for transcripts annotated with enriched GO terms, the relationship between length and association with RIN. Blue and red points represent genes annotated with the indicated GO term: genes we define as associated with RIN are highlighted in blue, and genes that fall below the cut-off used to define RIN-associated mRNAs are highlighted in red. Dashed purple horizontal lines indicate the median transcript length for all genes annotated with a given GO term. Solid vertical lines indicate no enrichment in the RIN RIP, and solid dark-grey horizontal lines indicate the median transcript length of all transcripts represented on the plot.

Together, these analyses lead to the hypothesis that RIN plays a global role in binding and potentiating the stability and translation of mRNAs that encode core components of gene expression and mitochondria in early embryos. Our analyses below of *rin*-mutant ovaries as well as our tethering studies in S2 cells, suggest that RIN/G3BP may play a similarly positive post-transcriptional role in additional cell types.

### Levels of RIN’s target transcripts are reduced in *rin* mutants

To assess the consequences of removal of endogenous RIN on the expression of RIN’s target mRNAs, we produced females trans-heterozygous for two previously identified *rin* alleles, *rin^2^* (Pazman et al., 2000) and *rin^3^* (Costa et al., 2013) (i.e., of genotype *rin^2^/rin^3^*) and used as controls females that were either *rin^2^/+* or *rin^3^/+*. We selected for analysis thirteen mRNAs that were highly enriched in the RIP-chip experiments, five that were unchanged, and seven that were depleted (Table S9). We used RT-qPCR to analyze transcript levels in at least three biological replicates from ovaries rather than embryos in order to exclude secondary effects in embryos from the mutant (Costa et al., 2013). Expression was normalized to the average of the transcripts with no enrichment for RIN association and the change in expression of the depleted, non-enriched, and enriched groups was compared pair-wise between *rin^2^/rin^3^* mutant ovaries and control ovaries using the two-tailed Wilcoxon signed-rank test. Transcripts depleted for RIN binding and transcripts non-enriched for RIN binding did not show a significant change in expression in mutant ovaries compared to control (z = 0.82, *P* = 0.41 and z = 0.12, *P* = 0.90, respectively). In contrast, transcripts enriched for RIN-binding showed a significant decrease in expression levels in mutant ovaries relative to control (z = 3.63, *P* < 3 x 10^-4^) (Table S9). These data are consistent with the hypothesis that RIN acts as a potentiator of the stability of its maternal mRNA targets.

### RIN and G3BP potentiate mRNA stability and translation in S2 tissue culture cells

Finally, to experimentally test the prediction that RIN is a potentiator of mRNA stability and translation, we tethered FLAG-tagged full-length RIN protein fused to BIV-Tat to a luciferase reporter mRNA containing six tandem BIV-TAR elements in its 3’UTR (Wakiyama et al., 2012) in *Drosophila* S2 tissue culture cells (Figure 6A). BIV-Tat-FLAG-RIN behaved the same way as endogenous RIN (Aguilera-Gomez et al., 2017): under non-stress conditions it was ubiquitously distributed in the cytoplasm and co-localized with CAPR (Figure S15). The fact that BIV-Tat-FLAG-RIN was ubiquitously distributed in the cytoplasm of unstressed cells is consistent with a report that overexpression of RIN in *Drosophila* S2 cells does not induce SGs (Aguilera-Gomez et al., 2017), and permitted us to assess the role of RIN in regulating target mRNA expression outside SGs.

**Figure 6.**
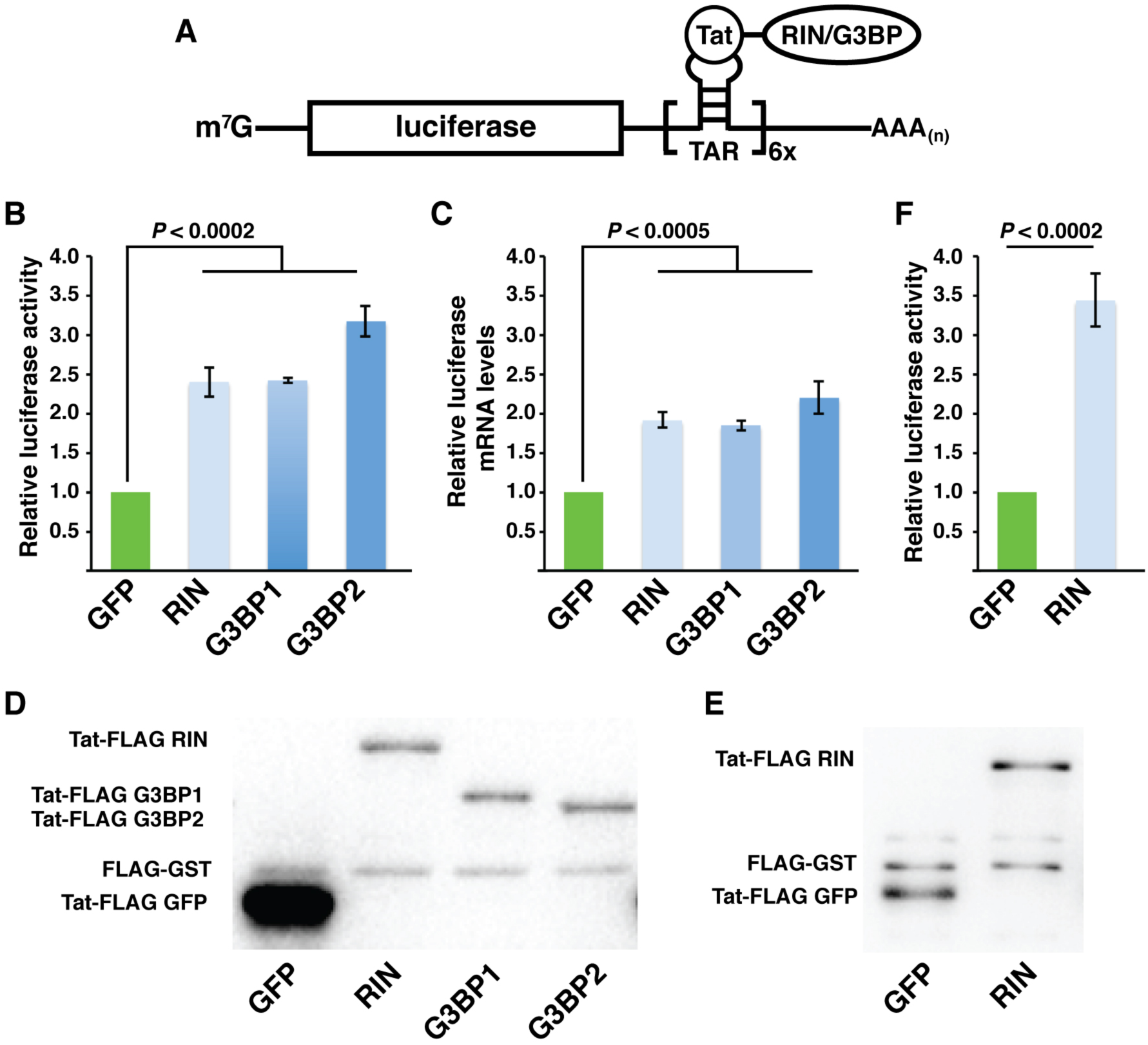
Tethering RIN or its human homologues G3BP1 or G3BP2 potentiates reporter expression. **(A)** A schematic representation of the tethering assay used to assess an RNA-binding protein’s function. The trans-activator peptide (Tat) interacts with stem-loop structures known as trans-activation response (TAR) elements. By co-expressing a luciferase reporter mRNA carrying six TAR elements in the mRNA’s 3’UTR, and RIN/G3BP-tagged Tat, RIN/G3BP will be tethered to the luciferase mRNA, thus allowing one to assess RIN/G3BP’s ability to control mRNA expression. **(B)** The Tat-tagged proteins indicated on the x-axis were co-expressed in cells with TAR-tagged firefly luciferase mRNA. The y-axis indicates the firefly luciferase enzyme activities normalized to a co-transfected unregulated *Renilla* luciferase. The normalized firefly luciferase enzyme activities for tethered RIN, G3BP1, G3BP2 were compared to tethered GFP, whose normalized luciferase activity was set to 1 (n=3). **(C)** The relative firefly luciferase mRNA levels were assessed by RT-qPCR, when the indicated protein on the x-axis was tethered. Firefly luciferase mRNA levels were normalized to *Renilla* luciferase mRNA levels and normalized GFP levels were set to 1 (n = 3). **(D)** Tethered proteins carry a triple FLAG, allowing us to compare their levels using anti-FLAG western blots. GST carrying a triple FLAG tag served as a loading/transfection control. **(E, F)** The same as **(D)** and **(B)**, respectively, with the exception that the levels of transfected Tat-GFP were reduced 10-fold. In all graphs, error bars indicate standard deviation and results of Student’s *t*-tests are indicated.

Tethering RIN resulted in a ∼2.4-fold increase in luciferase protein levels compared to a negative control in which a BIV-Tat-FLAG-GFP fusion protein was tethered to the same reporter (Figure 6B). To determine whether this increase in luciferase levels might be a result of increased translation, increased mRNA levels, or both, we also assayed luciferase mRNA levels. This revealed that RIN tethering resulted in a 1.9-fold increase in mRNA levels compared to the control (Figure 6C). These data show that RIN stabilizes the reporter mRNA and, because the relative increase in luciferase protein levels was greater than the relative increase in luciferase mRNA levels, are also consistent with a role for RIN in stimulation of translation.

We next assayed the expression levels of Tat-tagged RIN and GFP via western blot and found that GFP was expressed at significantly higher levels (Figure 6D). To assess whether the differential behavior of the TAR reporter in the presence of tethered RIN versus GFP could be explained by this difference in protein levels, we reduced the level of GFP plasmid transfected into cells, while keeping the levels of the RIN plasmid constant. This reduced the expression of GFP to levels equivalent to those seen for RIN (Figure 6E). When we compared luciferase expression in this experiment, the fold enhancement in luciferase expression mediated by tethered RIN was 3.4-fold (Figure 6F), confirming that tethered RIN enhances reporter expression.

To determine whether this might represent a conserved function of RIN, we tested the effect of tethering human G3BP1 or G3BP2 to the reporter mRNA in S2 cells. As was the case for BIV-Tat-FLAG-RIN, BIV-Tat-FLAG-G3BPs were ubiquitously distributed in the cytoplasm (Figure S15). Tethering either G3BP1 or G3BP2 led to increased levels of both luciferase protein and luciferase mRNA similar to those observed for RIN (Figure 6B, C). Thus, potentiation of mRNA stability and translation represent conserved functions of the RIN/G3BP family of proteins in unstressed cells.

## DISCUSSION

We have used a combination of IP-MS, polysome gradients, RIP-chip, computational analyses, RT-qPCR in mutants, and tethering experiments in tissue culture cells, to define RIN’s protein partners and bound mRNAs, and to elucidate RIN’s role as a post-transcriptional regulator in embryos and unstressed tissue-culture cells. Together our experiments have shown that *Drosophila* RIN and human G3BPs function to potentiate the stability and translation of bound mRNAs, contrasting with the repressive role of these RBPs in stressed cells.

### Features of RIN-bound transcripts

Two features, short transcript length and the motif bound by the RIN RRM *in vitro*, are predictive of RIN binding. That short transcripts in general might be more likely to contain the motif, is excluded by the fact that RIN’s target mRNAs are enriched for the motif when compared to length-matched, co-expressed, unbound mRNAs. Thus, we propose that each feature separately contributes to RIN target mRNA binding, with shortness playing a greater role than the motif (AUROC = 0.83 and 0.55 - 0.61, respectively).

While it is unclear how RIN is able to measure transcript length, there is precedent for the differential behaviour/regulation of short versus long mRNAs. For example, in general short mRNAs are more highly translated than long mRNAs and this is thought to reflect the fact that short mRNAs have a higher affinity for the cap-binding complex (Costello et al., 2015; Thompson and Gilbert, 2017). It has been proposed that this higher affinity is related to the possibility that short mRNAs are able to form a closed-loop structure more readily than longer mRNAs. We note that a recent study that calls the closed-loop model into question was unable to test short mRNAs because of technical limitations (Adivarahan et al., 2018). Thus, even if long mRNAs do not form a stable closed loop, it remains possible that short mRNAs do.

The ribosome-associated protein, RACK1, has been shown to be required for the efficient translation of short but not long mRNAs (Thompson et al., 2016). Although we have shown here that RIN is associated with polysomes in early embryos, *Drosophila* RACK1 is not on our list of RIN protein interactors, neither is there a significant overlap between our RIN protein interaction network and a set of protein interactors previously identified for RACK1 (Kuhn et al., 2017). Thus, the mechanisms by which RIN recognizes short mRNAs may differ from those that have been identified previously.

Another striking feature of RIN-bound mRNAs is that they are depleted for SREs, the binding sites of SMG. SMG destabilizes and translationally represses its target mRNAs. This could suggest that the high translational efficiencies and stability of RIN-target mRNAs is simply a result of a lack of SREs. However, we have provided evidence that RIN can exert its effects in situations where SMG protein is not present (i.e., during oogenesis, in embryos older than 3 hours, and in S2 cells). Thus we conclude that most RIN-bound mRNAs in the early embryo are upregulated directly by RIN and indirectly through a lack of SREs.

### RIN as a direct and indirect potentiator of gene expression

Since the target mRNAs of RIN are enriched for GO terms related to multiple levels of the core gene expression machinery –– transcription, splicing, and translation –– RIN may directly potentiate expression of its bound target transcripts and, in so doing, also indirectly upregulate gene expression globally. As an example of how RIN/G3BP’s direct and indirect effects might converge on the same cellular process, we consider the production of cytoplasmic ribosomes. Metabolic labelling with radioactive amino acids has shown that cytoplasmic ribosomal protein (cRP) synthesis increases after fertilization, peaks at 3-4 hours, and subsequently decreases (Santon and Pellegrini, 1980, 1981). Recent ribosome footprinting-based measurements have confirmed that the translational efficiency of cRP mRNAs increases in early embryos relative to mature oocytes (Eichhorn et al., 2016). We have shown here that RIN potentiates target mRNA stability and translation and that cRP mRNAs are highly enriched among RIN’s targets; thus, potentiation of cRP mRNAs is expected to be a direct effect of RIN.

cRP mRNAs are also regulated by their conserved 5’-terminal <u>o</u>ligo<u>p</u>yrimidine (TOP) motifs (Meyuhas and Kahan, 2015). Two RBPs, La and Larp1, have been implicated in regulation of the stability and/or translation of 5’TOP mRNAs via this motif (Cardinali et al., 2003; Crosio et al., 2000; Fonseca et al., 2018; Pellizzoni et al., 1996; Tcherkezian et al., 2014). We have shown here that the *Drosophila* orthologs of both of these RBPs––LA and LARP–– associate with RIN in an RNA-dependent manner. This could reflect the fact that LA, LARP and RIN co-bind cRP mRNAs and, thus, that this class of mRNAs is subject to multiple direct mechanisms that potentiate its expression. Noteworthy is the fact that, of our RIN target mRNAs, only the cRP transcripts carry 5’TOP motifs and, as such, these represent a distinct class of mRNAs.

A possible indirect role of RIN in regulation of cRP mRNAs relates to their upregulation by the Ragulator complex (Damgaard and Lykke-Andersen, 2011; Wilbertz et al., 2019). We have shown here that RIN binds mRNAs encoding all five subunits of the Ragulator complex. Thus, positive regulation by RIN of Ragulator subunit synthesis could indirectly promote cRP mRNA translation.

Direct and indirect regulation of other aspects of gene expression by RIN are also likely. Potentiation of production of core components of the transcription and splicing apparatus, as well as of the translation machinery, might ensure that none of these becomes rate-limiting for gene expression in rapidly developing early embryos. Likewise, adequate ATP production would be ensured by potentiation of expression of mitochondrial ribosomal proteins and components of the electron transport chain.

That said, our data suggest that the direct effects of RIN are much more pronounced than its indirect effects. Specifically, we have shown that, in *rin* mutants, levels of several target mRNAs are significantly reduced whereas co-expressed non-targets do not change significantly. It should be noted however, that these analyses were of a small subset of targets; global analyses –– which fall outside the scope of the current study –– might reveal that indirect targets also change significantly.

### A new function for RIN/G3BP sequestration into stress granules

Our data have important implications for our understanding of how cells respond to stress and the role of SGs in that response. A theme in the cellular stress response is a general downregulation of mRNA expression. For example, stress triggers eIF2 αphosphorylation, which prevents translation initiation (reviewed in Panas et al., 2016). This in turn triggers polysome disassembly, resulting in SG assembly. The storage of long mRNAs in SGs serves as a further mechanism to downregulate their translation. Based on our results we propose that the recruitment of RIN/G3BPs into SGs would sequester these proteins from their short target mRNAs in the cytoplasm, serving to downregulate the expression of these transcripts. This could indirectly downregulate global gene expression by limiting the production of proteins involved in translation, transcription and splicing, Likewise, if RIN/G3BPs serve to upregulate mitochondrial function and ATP production, sequestration could attenuate this aspect of cellular metabolism in stressed cells.

## Acknowledgements

We thank the three anonymous reviewers for helpful feedback regarding the manuscript, and the following for providing antibodies: Liz Gavis (anti-RIN) (Aguilera-Gomez et al., 2017) and Paul Macdonald (anti-CAPR) (Papoulas et al., 2010). The anti-FMR1antibody (5A11) developed by H. Siomi was obtained from the Developmental Studies Hybridoma Bank, created by the NICHD of the NIH and maintained at The University of Iowa, Department of Biology, Iowa City, IA 52242. During the course of this research, we made extensive use of FlyBase and data from the Berkeley *Drosophila* Genome Project. This research was supported by grants from the Natural Sciences and Engineering Research Council Discovery Grants to H.D.L. (RGPIN-06246) and to C.A.S. (RGPIN-435985), and from the Canadian Institutes for Health Research to Q.M. (MOP-125894).

## Author contributions

Conceptualization: J.D.L, C.A.S. and H.D.L.; Methodology: J.D.L., C.A.S., H.D.L., S.A., Q.M., S.S.S and J.T.W.; Investigation: J.D.L., J.L., A.K.W., S.L., A.K., G.B., W-X.C., N.J-L. and A.K.; Software and Formal Analysis: K.L., J.D.L. and Q.M.; Writing –– Original Draft: J.D.L., J.L.; Writing –– Review and Editing: H.D.L. and C.A.S.; Funding Acquisition: H.D.L., C.A.S and Q.M.; Resources: S.A., S.S.S. and J.T.W; Supervision: C.A.S., H.D.L., S.A. and Q.M.

## STAR METHODS

### CONTACT FOR REAGENT AND RESOURCE SHARING

Further information and requests for resources and reagents should be directed to and will be fulfilled by the Lead Contact, Howard Lipshitz.

### EXPERIMENTAL MODEL AND SUBJECT DETAILS

Wild-type *Drosophila* stock was *w^1118^*; mutant lines were *rin^2^*/*TM6B, Sb* (BDSC# 9303) (Pazman et al., 2000), and *rin^3^*/*TM6B, Sb Tb* (BDSC# 57694) (Costa et al., 2013). The latter two lines were crossed to produce *rin^2^/rin^3^* mutants for RT-qPCR analysis. *Drosophila* S2 tissue culture cells were maintained at 25°C in Express Five SFM (Fisher Scientific) containing 100units/mL penicillin, 100µg /mL streptomycin and 16mM glutamine.

### METHOD DETAILS

#### Western blots

Embryos were collected from cages containing *w^1118^* flies, dechorionated with 100% bleach for 2 minutes, washed with 0.1% Triton X-100 and lysed by crushing in a minimal volume of lysis buffer: 150 mM KCl, 20 mM HEPES-KOH pH 7.4, 1 mM MgCl_2_, supplemented with protease inhibitors (1 mM AEBSF, 2 mM benzamidine, 2 µg/mL pepstatin, 2 µg/mL leupeptin) and 1 mM DTT. The lysate was cleared by centrifugation (15 min at 4°C, 13000 RPM) and stored at –80°C. Protein concentration was determined using the Bio-Rad Protein Assay Dye Reagent (Cat#5000006) and 15µg total protein was resolved by SDS-PAGE. For S2 cell western blots, 1.2×10^6^ cells were pelleted via centrifugation and lysed with 2xSDS sample buffer supplemented with 1 mM AEBSF and boiling for 3 minutes. Two to eight μL of the resulting extracts were resolved via SDS-PAGE. Proteins were transferred to PVDF membrane, blocked at room temperature for 1 hour with 0.5% milk in PBST (1x PBS + 0.1% Triton X-100). Blots were then incubated with the appropriate antibodies: anti-RIN (1:50000) (Aguilera-Gomez et al., 2017) and anti-α tubulin (1:10000, Sigma-Aldrich T5168), or anti-FLAG (1 μg/mL, Sigma-Aldrich F3165) at 4°C overnight. After incubation with primary antibody, the blot was incubated with HRP-conjugated secondary antibody (1:5000; either of Peroxidase-Affinipure goat α-mouse IgG (H+L) Cat#115-035-003 or goat α-rabbit IgG (H+L) Cat#115-035-144, Jackson Immunoresearch) at room temperature for 1 hour. Western blots were developed using the ECL detection system (Millipore Immobilon Luminata Crescendo Western HRP substrate Cat#WBLUR0500). Relative levels of RIN and α-tubulin were determined using a standard curve. Western blots were imaged and quantified using ImageLab (BioRad).

#### Immunostaining of embryos

Standard immunostaining procedures were followed (Ashburner, 1989). Embryos were collected after a 4 hour egg-lay from cages containing either *w^1118^*, *rin^2^/rin^3^* mutant females or females heterozygous for *rin*. Embryos were dechorionated with 50% bleach and fixed in formaldehyde (4%) and methanol. To visualize RIN and FMR1, fixed embryos were incubated with the following primary antibodies used at the concentrations indicated: rabbit anti-RIN (1:1000; provided by Liz Gavis) (Aguilera-Gomez et al., 2017) and mouse anti-FMR1 (1:10; #5A11 from the Developmental Studies Hybridoma Bank) (Okamura et al., 2004). Conjugated secondary antibodies were purchased from Invitrogen-ThermoFisher and used at 1:300 (Alexa Fluor555 Goat Anti-Rabbit IgG, Catalog number A21429, and Alexa Fluor488 Goat Anti-Mouse IgG, Catalog number A11029). Embryos were also labeled for DNA using a 0.001mg/ml DAPI (Sigma, Catalog number D9542) incubation for 10 min. Images were collected using a Nikon Ti-S inverted microscope with Nikon C2 confocal system, using NIS Elements AR software. Images were then processed for figures using Fiji/ImageJ (Schindelin et al., 2012; Schneider et al., 2012), Adobe Photoshop and Adobe Illustrator.

#### Generation of anti-RIN synthetic antibodies

Synthetic antibodies were generated against antigens comprising RIN amino acids 12-131 (antibodies D072) or amino acids 12-170 (antibody D074; all amino acid numbering according to RIN-PB isoform), which were expressed and purified from *E. coli* as GST fusion proteins as described (Laver et al., 2012). Synthetic antibodies were obtained by performing five rounds of binding selection with synthetic antibody Library F (Persson et al., 2013) against each of the two RIN antigens, as described (Laver et al., 2015a). The three synthetic antibodies obtained against RIN were expressed and purified from *E. coli* as Fabs, tagged at the C-terminus of the light chain with a FLAG tag, as described (Laver et al., 2012).

#### Immunoprecipitations and mass spectrometry

For RIN IP-MS experiments, 0–3 h old embryos were collected, and lysed by crushing in a minimal volume of lysis buffer (150 mM KCl, 20 mM HEPES-KOH pH 7.4, 1 mM MgCl_2_, supplemented with protease inhibitors and 1 mM DTT), followed with clearing by centrifugation (15 min at 4°C; 20,000 x g), and stored at –80°C. Immediately prior to performing IPs, this cleared lysate was thawed and diluted ½ with lysis buffer, and supplemented with Triton X-100 to a final concentration of 0.1%. For IPs, ∼800 µL of this diluted lysate, with or without 0.35 µg/µL RNase A, was incubated with 40 µL of anti-FLAG M2 beads (Sigma) that were pre-loaded with 20 µg of either RIN Fab D072 or control C1 Fab, and blocked with BSA. IPs were incubated for ∼3 hours at 4°C with end-over-end rotation. After incubation, beads were washed 4–5 times with lysis buffer supplemented with 0.1% Triton X-100, twice with lysis buffer (no Triton X-100), then transferred to new tubes and washed twice with lysis buffer (no Triton X-100). Bound proteins were eluted by tryptic digest: beads were resuspended in 200 µL of 50 mM ammonium bicarbonate pH 8, supplemented with 2 µg of trypsin, and incubated overnight at room temperature with end-over-end rotation. The following day, the digested supernatant was recovered, and beads were washed once with an additional 200 µL of 50 mM ammonium bicarbonate to collect any residual eluted material. The two supernatants were pooled and dried by speed-vac. Liquid chromatography–tandem mass spectrometry (LC-MS/MS) was performed using the Thermo Q-Exactive HF quadrupole-Orbitrap mass spectrometer (Thermo Scientific) and the methods followed those previously described (Chiu et al., 2016; Jiang et al., 2015; Liu et al., 2014).

Three biological replicates of RIN IPs and control IPs were performed, in both the presence and absence of RNase A. Data analysis is described below.

#### Polysome gradients

Embryos were collected 0-3 hours post egg laying and lysed in a 2 mLs of lysis buffer per gram of embryos. Lysis buffer was 50 mM Tris pH 7.5, 2mM MgCl_2_, 150 mM KCl, 100 µM GTP, 1 mM DTT, 50 U/mL RNase inhibitor, 1 mM AEBSF, 2 µg/mL leupeptin, 2 mM benzamidine, 2 µg/mL pepstatin A that was supplemented with either 0.5 mg/mL cycloheximide or 2mM puromycin. After lysis samples were left on ice for 20 minutes and then incubated at 30 °C for 10 minutes. 30 % Triton-X100 was added to a final concentration of 1% and samples were spun at 6000 x g for 10 minutes. 400 µL the resulting supernatant was layered onto a 5mL 15% to 45% sucrose gradient in 7.5 mM MgCl_2_, 500 mM NaCl, and 50 mM Tris pH 7.5. The gradient was created using a BioComp Model 117 Gradient Mate gradient maker (BioComp) using according to the manufacter’s instructions. The gradients were chilled on ice for 1 hour before extract application after which they were spin at 36,000 rpm for 2h 30 minutes at 4 °C in a Beckman SW50.1 rotor. The gradients were then hand fractionated into nine 600 µL fractions which were analyzed via western blot.

#### RNA co-immunoprecipitations

For RNA co-immunoprecipitations for RIP-Chip experiments, immunoprecipitations were carried out from 400 µL of embryo lysate, prepared from embryos collected 0–3 h post-egg-laying, using 20 µg of FLAG-tagged Fab captured on 40 µL of anti-FLAG M2 affinity gel (Sigma), as previously described (Laver et al., 2015a). RNA co-immunoprecipitations for RT-qPCR experiments were performed similarly but at a reduced scale, using one-eighth the amount of material listed above.

#### Microarray analysis of RIN RIP samples

Microarrays were custom-designed *Drosophila* 4 x 72 K NimbleGen arrays (GEO platform number: GPL10539). RIP samples were reverse-transcribed, labelled, and hybridized as previously described (Laver et al., 2015a). Three biological replicates each were performed for RIN and control immunoprecipitated samples. Arrays were scanned, quantified, and normalized as previously described (Laver et al., 2015a), with all RIN and control IP samples normalized together. The data have been deposited in the Gene Expression Omnibus (GEO) under accession number GSE12900. Analysis is described below. [<u>For reviewers</u>: To review GEO accession GSE129900:

Go to https://www.ncbi.nlm.nih.gov/geo/query/acc.cgi?acc=GSE129900 Enter token otwtmoeqjhcztmv into the box]

#### RT-qPCR

For RT-qPCR from RIN RNA co-immunoprecipitations, single-stranded cDNA was synthesized by reverse transcription of immunoprecipitated RNA with Superscript IV reverse transcriptase (Invitrogen) using a mixture of random hexamer and anchored oligo-dT primers, as described for cDNA synthesis for microarray sample preparation (Laver et al., 2015a). The single-stranded cDNA was used to perform quantitative real-time PCR with primers specific to the various transcripts assayed, using SensiFAST SYBR PCR mix (Bioline) and a CFX384 Real-Time System (Bio-Rad). Sequences of the primers used for PCR are listed in Table S10.

#### S2 cell transient transfection, dual-luciferase assay and RT-qPCR

*Drosophila* S2 tissue culture cells were maintained at 25°C in Express Five SFM (Fisher Scientific) containing 100units/mL penicillin, 100µg /mL streptomycin and 16mM glutamine. A mixture of 1.5ng Firefly luciferase-6x TAR plasmid, 1.5ng *Renilla* luciferase plasmid, 3ng Tat-FLAG-effector plasmid (carrying either the GFP, RIN, G3BP1 or G3BP2 open reading frame), 1.5ng of FLAG-GST and 192.5ng pSP72 was transfected into 0.4mL of S2 cells at a density of 1.75 x 10^6^ cells/mL using 0.4μL TransIT-Insect transfection reagent (Mirus Bio), according to the manufacturer’s instructions. Both luciferase reporters were cloned into pRmHa3 (Mohan et al., 2014) and, thus, were under the control of the metal-inducible metallothionein promoter. All FLAG-tagged constructs were derived from pAc5.1/V5-His (Thermo Fisher Scientific), which carries the Actin5C promoter. The expression of luciferase reporters was induced 24 hours post transfection through the addition of copper sulfate to a final concentration of 0.5mM. 24 hours post induction, Firefly and *Renilla* luciferase activities were measured using the Dual-Luciferase Reporter Assay System (Promega). In indicated experiments, the amount of transfected Tat-GFP was reduced to 0.3ng and 2.7ng of a pAc5.1/V5-His derivative expressing only Tat-FLAG was included to maintain equal levels of actin promoter across all transfections.

To assay reporter transcript levels, 48 hours post-transfection S2 cells were harvested, resuspended in TRI reagent (MRC) and RNA was purified according to the manufacturer’s protocol. 1ug of total RNA was treated with DNase I (Invitrogen) after which it was used to generate cDNA through reverse transcription with Superscript IV reverse transcriptase (Invitrogen) and random hexamers (Thermo Fisher) following the manufacturer’s instructions. The cDNA was subjected to quantitative real-time PCR using the SensiFAST SYBR PCR mix (Bioline) PCR mix and primers against the Firefly and *Renilla* luciferase ORFs. Relative levels of the Firefly and *Renilla* transcripts were determined using a standard curve. Sequences of the primers used for PCR are listed in Table S10.

#### S2 cell immunofluorescence and microscopy

Forty-eight hours post-transfection S2 cells were grown at 25°C on poly-D-lysine coated coverslips for 3 hours. Following any treatment, cells were immediately fixed with 4% EM grade formaldehyde in PBS for 10 minutes. Cells were rinsed once with PBSTx (1X PBS+0.1% Triton X-100), permeablized with PBSTx for 15 minutes then incubated with 5µg/mL α-FLAG M2 (F3165, Sigma) and 1:200 rabbit α-CAPR (Papoulas et al., 2010) overnight at 4C in a humidified chamber. Coverslips were washed 3 times for 5 minutes each time with PSBTx then incubated with α-mouse Alexa Flour 555 and α-rabbit Alexa Fluor 488 (Invitrogen-ThermoFisher Catalog numbers A21424 and A11034, respectively) for 2 hours at room temp in a humidified chamber. Coverslips were washed 3 times for 5 minutes each with PSBTx and then mounted onto slides with Fluoromount g + DAPI (00-4959-52, Invitrogen) and incubated overnight at 4°C. All images were collected using a Nikon Ti-S confocal microscope with NIS Elements AR software. Images were processed using ImageJ.

### QUANTIFICATION AND STATISTICAL ANALYSIS

#### Mass spectrometry

To identify RIN-interacting proteins, we used the ProHits software package (Liu et al., 2010) to perform Significance Analysis of INTeractome (SAINT), comparing RNase-treated RIN IP versus control IP samples, and non-RNase-treated RIN IP versus control IP samples. Specifically, SAINT input files were generated using the “TPP iProphet” search engine option in ProHits, filtering for iProphet probability > 0.95 and number of unique peptides < 2. SAINTexpress (exp3.3) was run using the ProHits interface, including only detected *Drosophila* proteins, with the following settings: number of compressed controls = 3; burn-in period, nburn = 2000; iterations, niter = 5000; lowMode = 1; minFold = 1; normalize = 1; nCompressBaits = 2. Identified proteins were defined as RNA-independent or RNA-dependent RIN-interacting proteins, if, in the analyses of the respective samples, they achieved a SAINT score ≥ 0.95 and a Bayesian false discovery rate (BFDR) ≤ 0.01.

#### Microarrays

To identify RIN-associated mRNAs, microarray data were analyzed using the Significance Analysis of Microarrays (SAM) (Tusher et al., 2001) function available in the MultiExperiment Viewer software application (Saeed et al., 2006; Saeed et al., 2003), as previously described (Laver et al., 2015a). Genes whose mRNAs were significantly enriched in the anti-RIN IPs compared to the control IPs, with an FDR of less than 5% and at least two-fold enrichment, were defined as RIN-associated mRNAs.

For the purposes of all subsequent analyses, FBgn gene IDs listed in the array annotation file were updated to FlyBase release 6.20 using the ‘Upload/Convert IDs’ tool available on FlyBase, and mRNAs corresponding to gene models which have since been withdrawn were excluded.

#### Extraction of length-matched target and non-target sets

Length-matched targets and non-targets were extracted as follows: first, we randomly chose a target transcript from the target set, and then we looked for a transcript in the non-target set that had the minimum length difference from that target, unless the length differences for the remaining non-targets were all higher or equal to the length of the target transcript (we randomly chose one if multiple non-targets with the same length difference were available). Non-targets were removed from the candidate non-target set when they were selected as a match for a target, and targets were removed from the candidate target set whether or not they had a match in the non-target set. We repeated this process until we exhausted the target set. Because RIN targets are generally shorter than non-targets, and the order of RIN targets being chosen from the target set was different for each matching process, the composition of the length-matched target and non-target sets was different in each iteration (five such iterations were carried out; see Figure S7). We extracted multiple length-matched target and non-target sets for 5′UTR, CDS, 3′UTR and full mRNA, using the sequence length of each transcriptomic region respectively.

#### Enrichment test of published RIN motifs

To test whether published RIN motifs are enriched in genes co-immunoprecipitated with RIN in our analysis, we used *in vitro*-determined *Drosophila* RIN and human G3BP2 motifs (Cook et al., 2011; Ray et al., 2013) (see http://cisbp-rna.ccbr.utoronto.ca/), and *in vivo* motifs of human G3BP1 and G3BP2 (Edupuganti et al., 2017). Position frequency matrices (PFMs) of the *in vivo* motifs were estimated from the height of base logos and produced using ggseqlogo (Wagih, 2017) and Logomaker (Tareen and Kinney, 2019). To eliminate the effect of sequence length on motif enrichment, we randomly selected corresponding transcript regions from length-paired co-expressed unbound genes as negative set for each transcript region of genes in the RIP-enriched gene sets (positive set). The longest transcript isoform of each gene was used. For a motif of length K and for all transcript subsequences of length K (i.e., K-mers), we calculated a “hit score” by multiplying the probability of that K-mer under the PFM and the probability of the entire subsequence being accessible at the given location within the transcript. The probability of a K-mer under a PFM was calculated by multiplying the probability of each base of the K-mer at its corresponding position in the PFM. Subsequence accessibility was estimated using RNAplfold (Bernhart et al., 2006; Lorenz et al., 2011) with the parameters: -W 80 -L 40 –u <motif_length>. For each motif, we tested its ability to distinguish positive genes from length-paired negative genes by ranking genes according to the maximum “hit score” of all subsequences of its corresponding transcript for that motif. We repeated the test 1000 times with length-paired negatives randomly selected each time, and reported the number of significant Mann-Whitney *U* tests, and the mean and standard deviation of the AUROCs and *P* values of each motif’s significant tests (Table S4). To show the distribution of motifs in RIN targets v.s. non-targets, we grouped the number of RIN motifs (defined as sites with “hit score” > 0.001) in a length-paired positive and negative sample from each of the three transcript regions into 20 bins, and used Gaussian kernels to fit the un-normalized positive and negative histogram for each region. The differences between the two corresponding Gaussian kernels in each of the three regions are displayed in Figure 2D.

#### Comparisons of RIN-associated mRNAs to published *Drosophila* datasets

For all comparisons of RIN-associated mRNAs to *Drosophila* transcripts reported in previously published datasets, all gene IDs from the various datasets were first updated using the FlyBase ‘Upload/Convert IDs’ tool to FBgn IDs from FlyBase release 6.20, to ensure consistency of identifiers between datasets. The lengths of the longest and shortest mRNA transcript isoforms for every gene were obtained from FlyBase release 6.20.

For comparisons of RIN-associated mRNAs to mRNA translation and abundance data from Eichhorn et al. (Eichhorn et al., 2016), only genes measured in both datasets were included in the scatterplot comparisons. In cases where a single gene had more than a single fold-enrichment value in our microarray data, due to the presence of multiple probe sets representing the same gene, the higher fold-enrichment value was used.

For comparisons of RIN-associated mRNAs to lists of genes from other datasets, including maternal decay classes, and N^6^-methyladenosine (m^6^A) modification, significant enrichment or depletion was assessed between the relevant lists with Fisher’s exact test, using as a background the intersection of the set of expressed genes for our RIP-Chip experiment and the set of genes included in the dataset being compared, as previously described (Laver et al., 2015a).

#### Comparisons of RIN-associated mRNAs and proteins to datasets in other species

To compare *Drosophila* RIN-associated mRNAs and proteins to reported lists of human G3BP-associated proteins, and stress granule enriched or depleted proteins and mRNAs in yeast, mouse, and human, the *Drosophila* homologs of genes in the non-*Drosophila* datasets were obtained using the DRSC Integrative Ortholog Prediction Tool (DIOPT) web server (http://www.flyrnai.org/cgi-bin/DRSC_orthologs.pl) (Hu et al., 2011). Only *Drosophila* homologs with a ‘Rank’ of ‘moderate’ or ‘high’ in the DIOPT output were considered, and where more than one *Drosophila* homolog existed, only the best match was included. After obtaining the lists of homologs, comparisons between the RIN-associated mRNAs and *Drosophila* homologs of datasets from other species were carried out as for the intra-species comparisons described above.

#### Gene ontology annotation enrichment analysis

GO annotation enrichment analysis was carried out using the DAVID 6.8 functional annotation tool web server (Huang da et al., 2009a; Huang da et al., 2009b), to search for enrichment of GO terms included in the GO FAT database. Genes identified by RIP-Chip as encoding RIN-associated mRNAs were analyzed for enrichment against the set of expressed genes defined for our RIP-Chip experiment, as previously described (Laver et al., 2015a). GO terms enriched at an FDR of less than 10% were considered significant.

For GO terms highlighted in scatterplots that depict transcript length and transcript translational status or abundance, lists of all genes annotated with each GO term were obtained from FlyBase using the controlled vocabulary search tool.

### DATA AND SOFTWARE AVAILABILITY

Microarray data have been deposited in the Gene Expression Omnibus (GEO) under accession number GSE12900. Reviewers, please go to https://www.ncbi.nlm.nih.gov/geo/query/acc.cgi?acc=GSE129900.

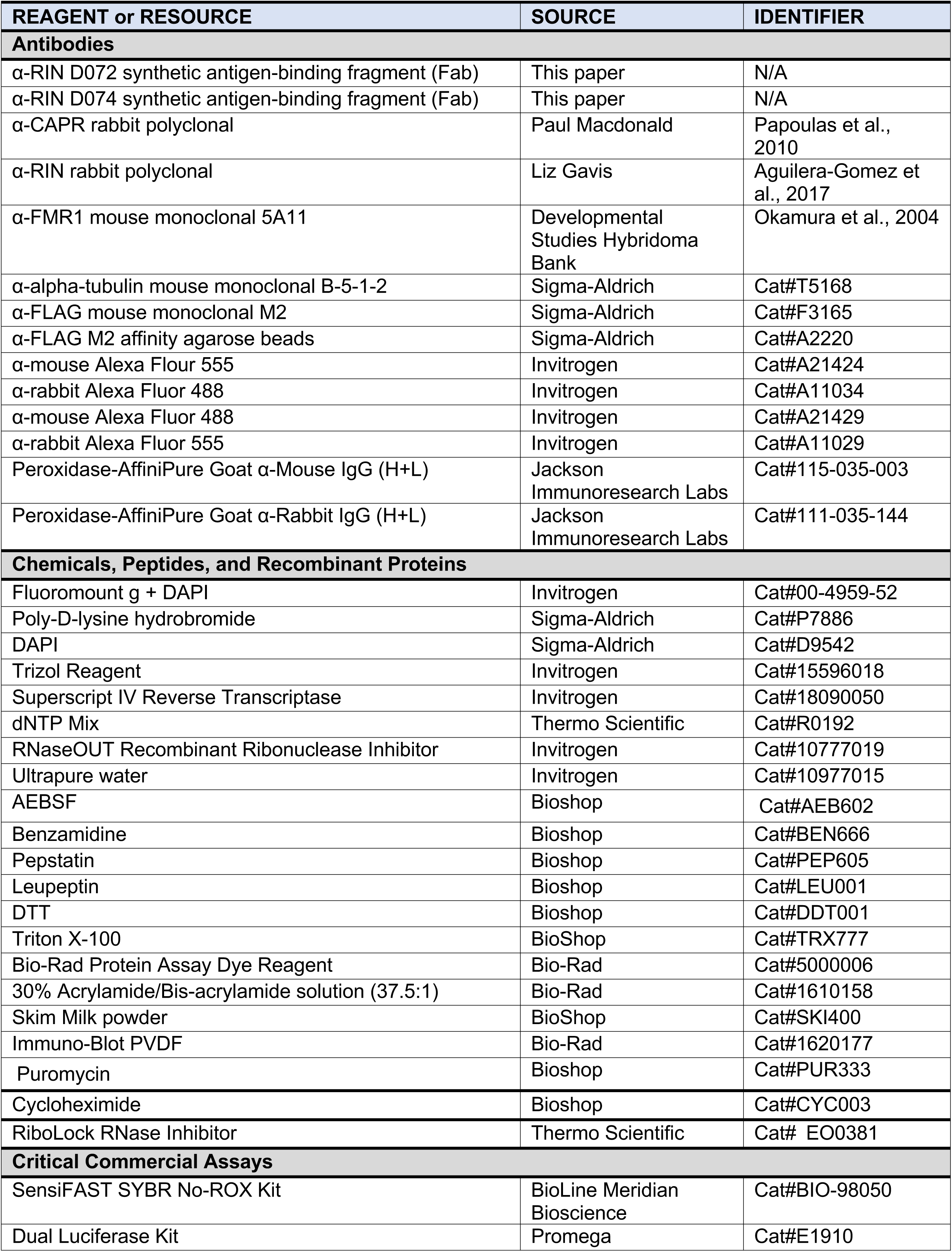
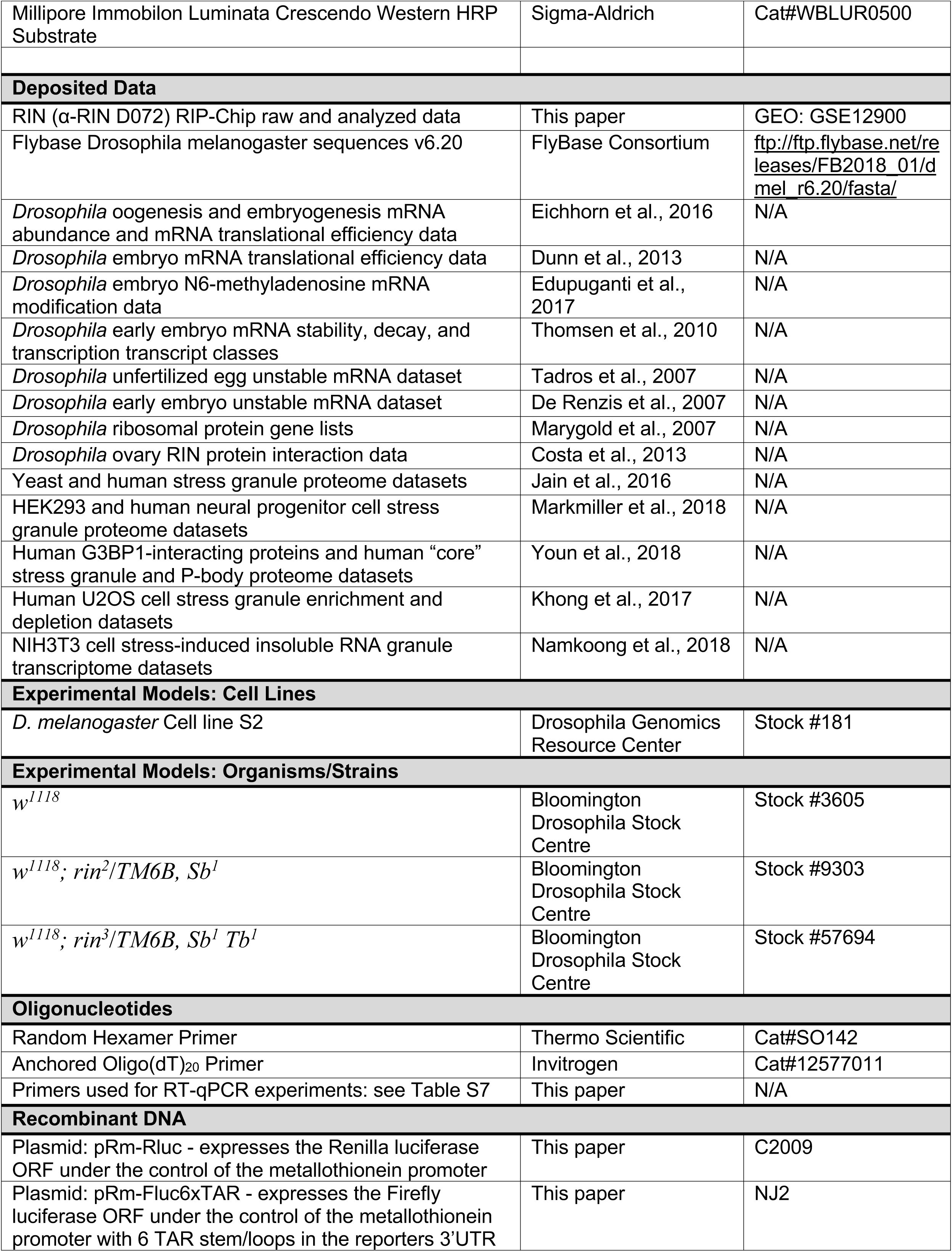
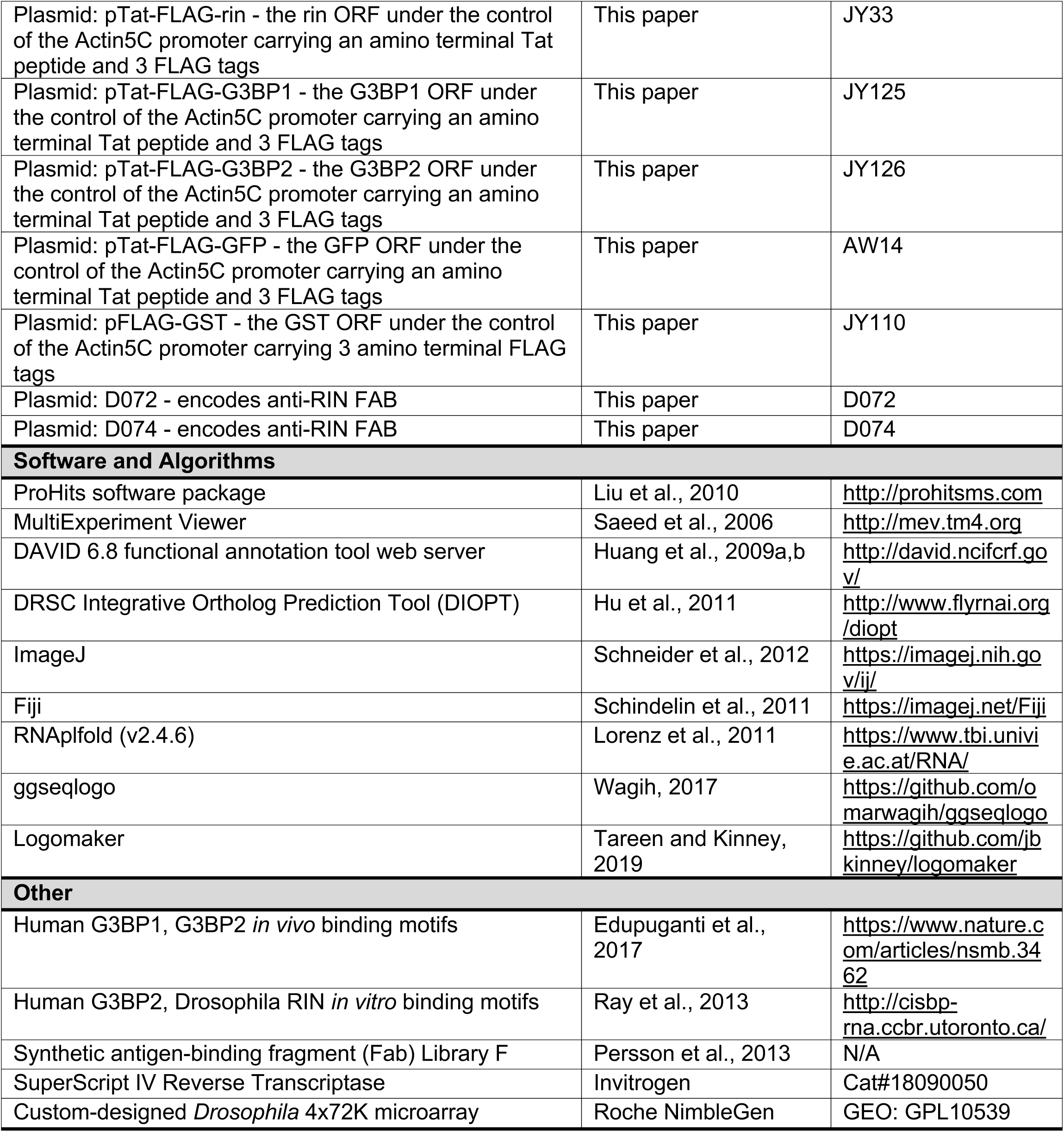

## List of Supplemental Figures and Tables with Legends

**Figure S1.**
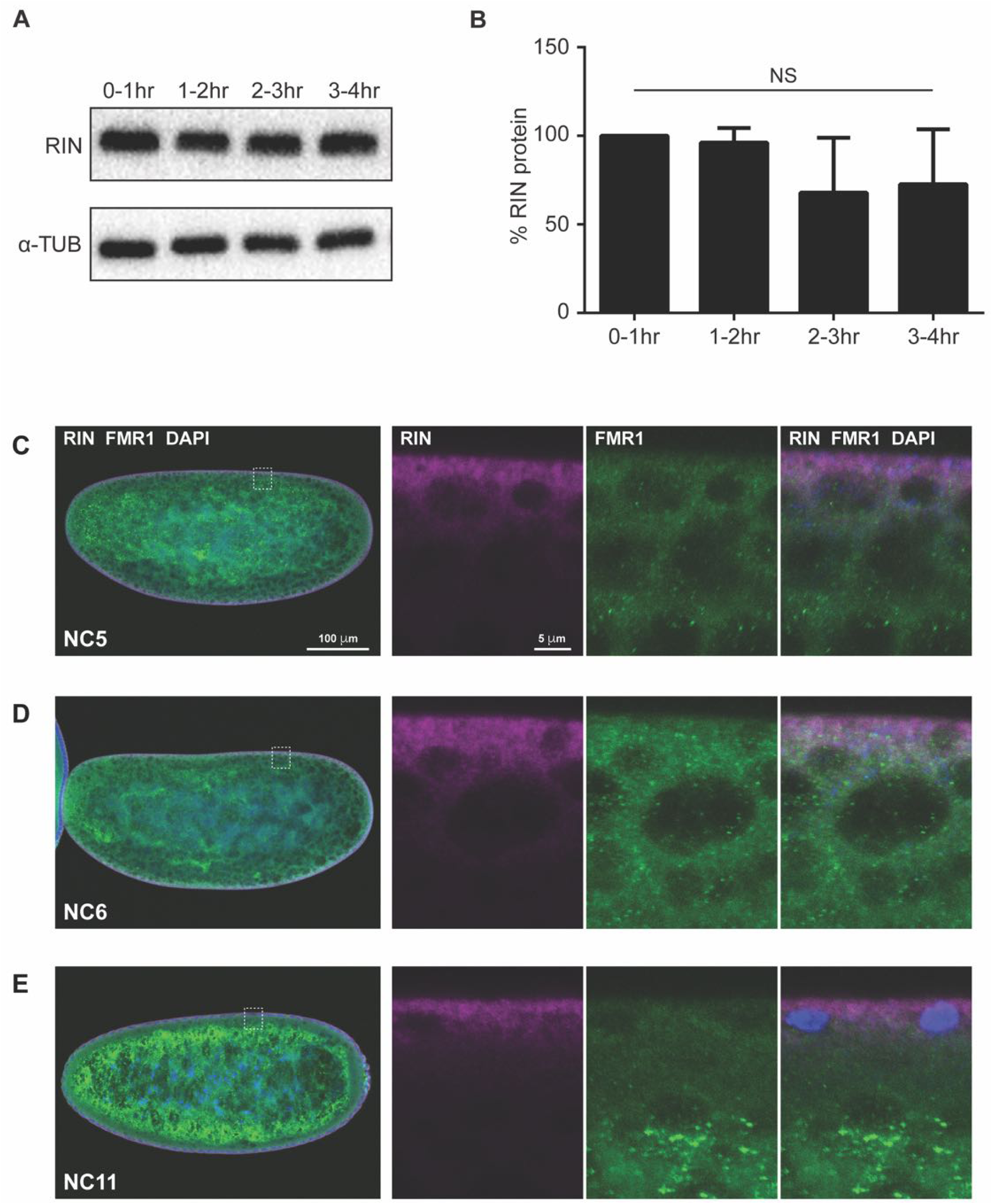
RIN expression in early embryos. **(A)** Western blots showing a time course of RIN expression during the first four hours of embryogenesis in *w^1118^* embryos. α-tubulin serves as the loading control. **(B)** RIN levels normalized to >-tubulin, with the 0–1 hr timepoint set to 100%. There is a slight but not statistically significant decrease of RIN levels between 1–2 and 2–3 hr (Student’s *t* test; error bars indicate standard deviation of three biological replicates). **(C)-(E)** Confocal immunofluorescence z-axis projections of three mid-sagittal sections of early embryos which show RIN (magenta) and FMR1 (green) at nuclear cycle 2 **(C)**, 6 **(D)** and 12 **(E)**. Nuclei were visualized with DAPI (blue). The scale bar in the left panel in (C) also refers to the equivalent panels in (D) and (E); the scale bar in the second-from-left panel in (C) also refers to the equivalent panels in (D) and (E). Controls for the specificity of the anti-RIN antibody are shown in Figure S2.

**Figure S2.**
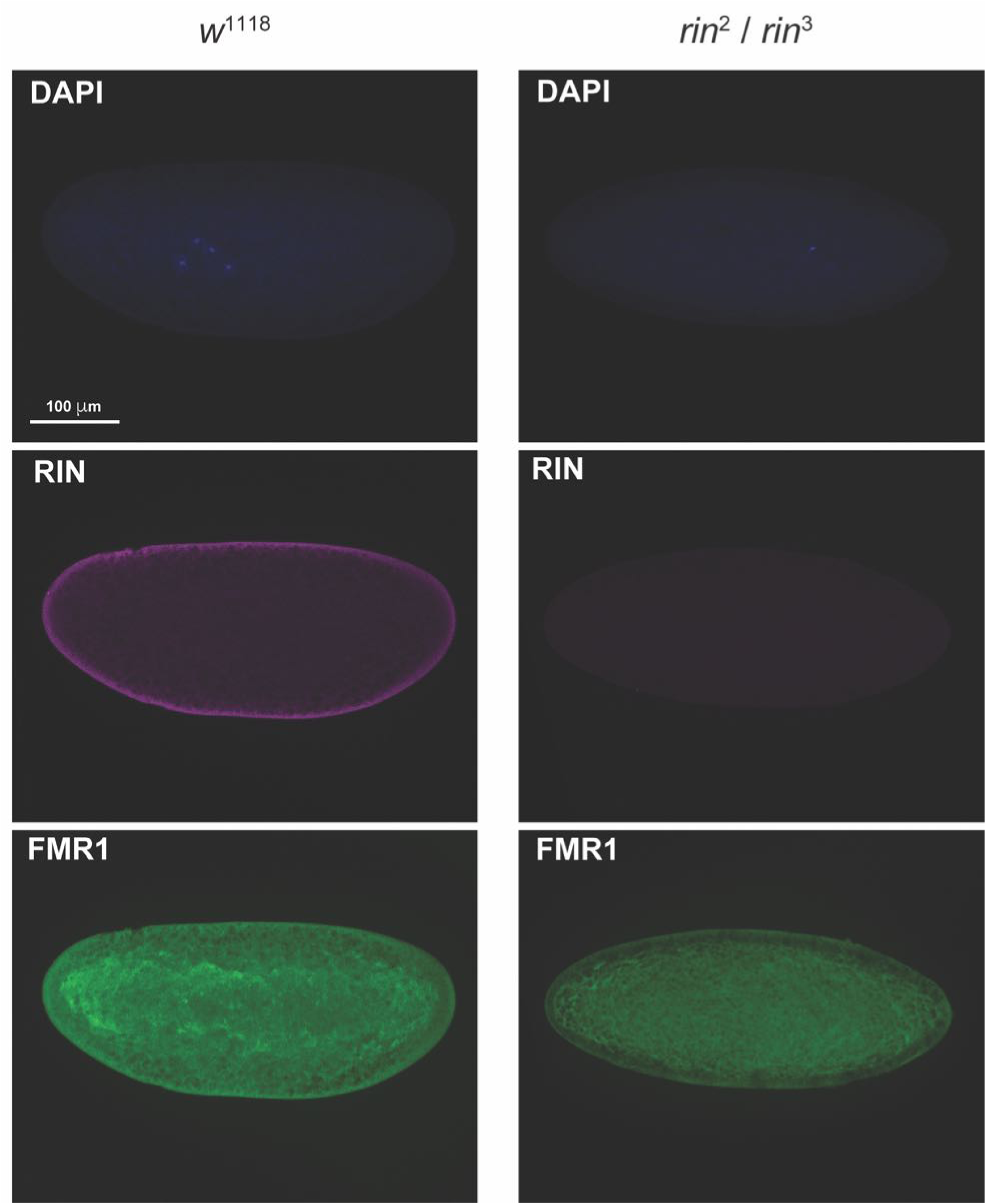
Specificity of the anti-RIN antibody for immunofluorescence. Shown are early embryos from ‘wild type’ *w^1118^* (left) or *rin^2^/rin^3^* mutant females. Confocal microscope z-axis projections of three mid-sagittal sections are presented. There is no RIN signal (magenta) in the mutants whereas control, FMR1, signal (green) is present in both genotypes.

**Figure S3.**
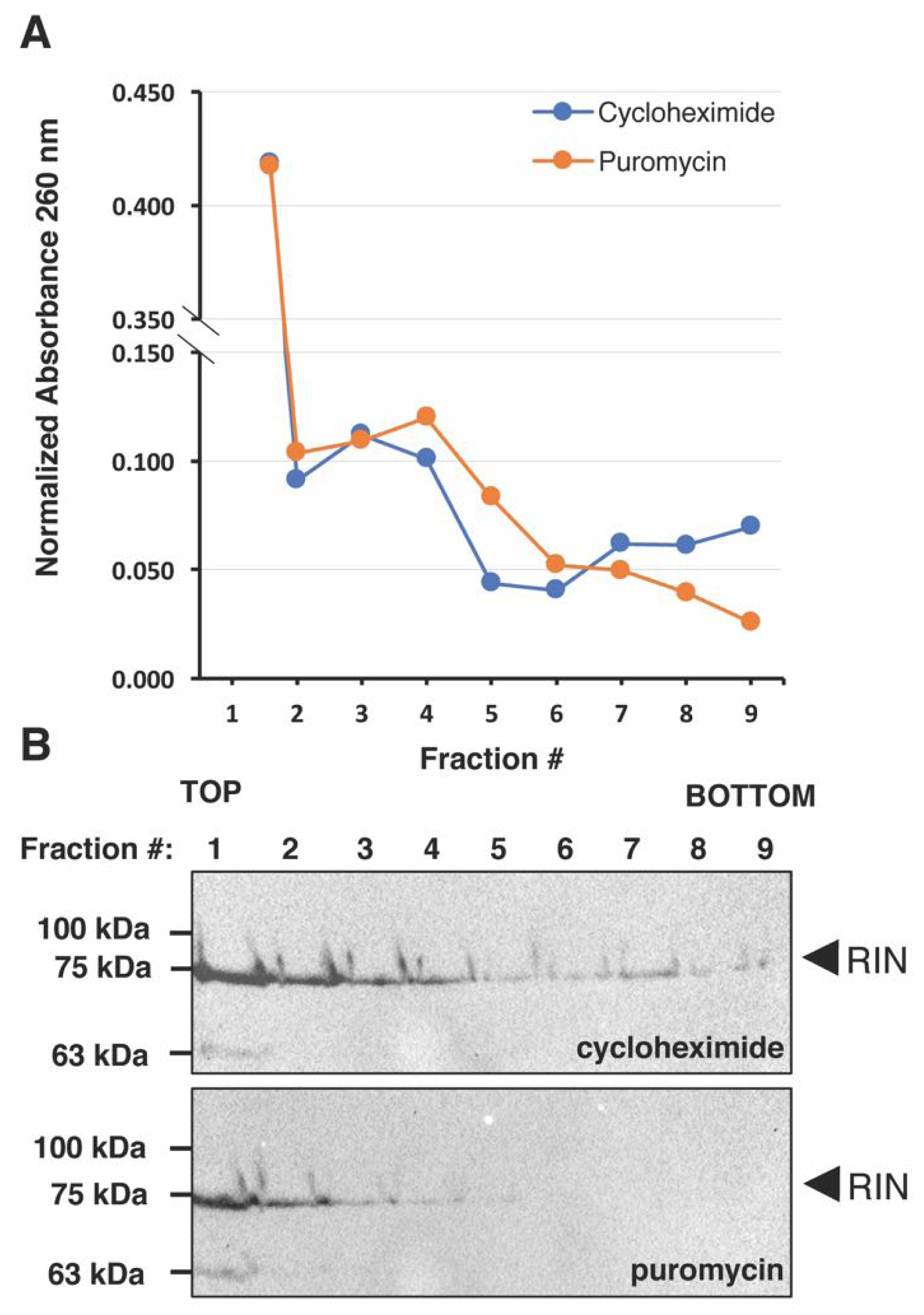
RIN is polysome associated. Embryo extract treated with either cycloheximide or puromycin was fractionated on sucrose gradients. **(A)** A_260_ of each fraction showing a decrease of absorbance in the denser fractions upon puromycin treatment. **(B)** Western blots for RIN.

**Figure S4.**
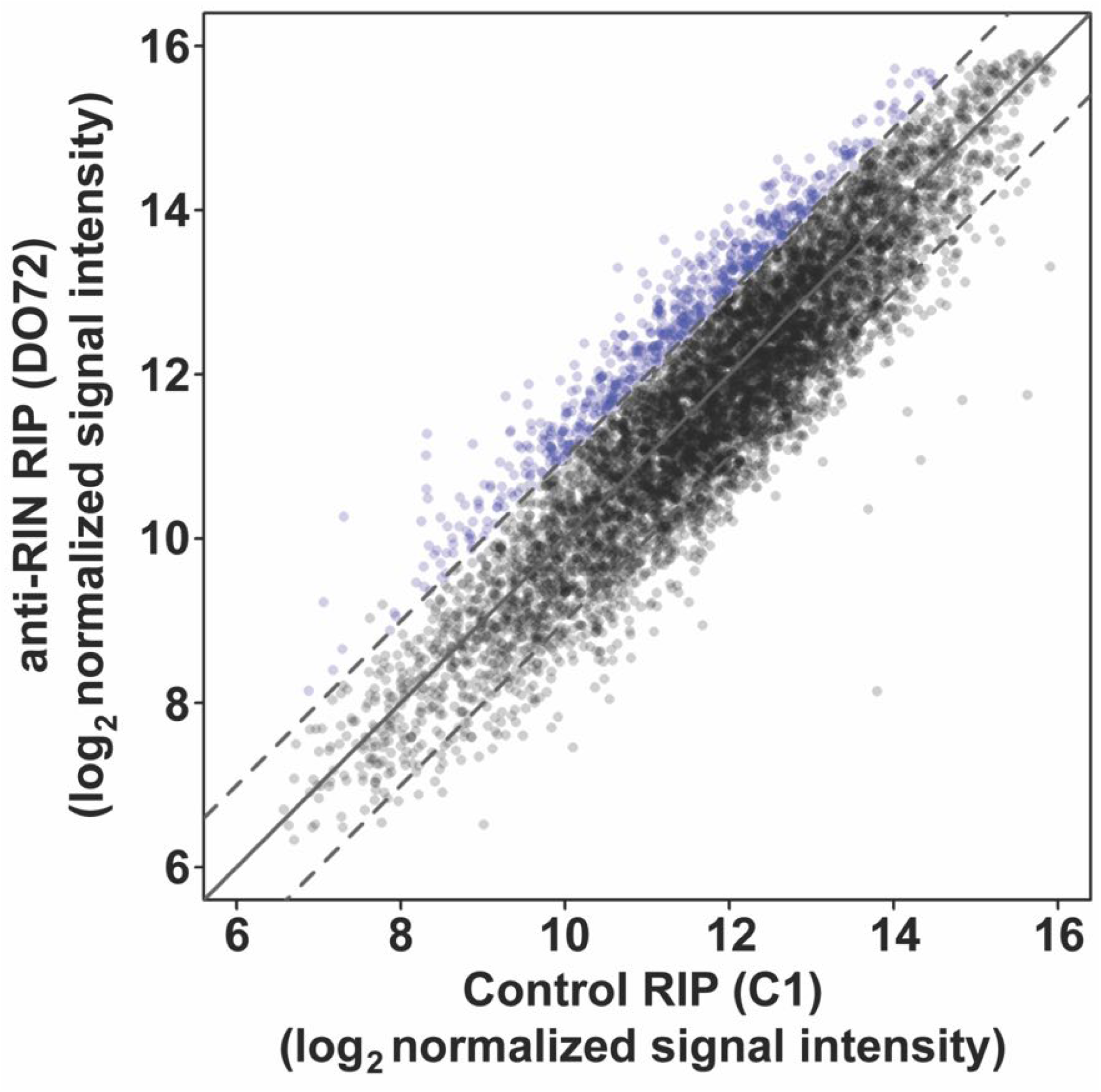
RIN associates with hundreds of mRNAs in early *Drosophila* embryos. Plot showing mRNA abundance, measured as the RMA-normalized microarray signal intensity, in anti-RIN (D072) RIPs versus C1 control RIPs, for all transcripts represented on the microarray that were defined as expressed in early embryos. Values represent the average of three biological replicates. mRNAs with an average enrichment > 2-fold in RIN vs C1 IPs with an FDR < 5% are defined as RIN-associated mRNAs and are highlighted in blue (606 transcripts corresponding to 566 genes). The solid diagonal line represents no enrichment, and dashed diagonal lines represent 2-fold enrichment or depletion. See also Table S3; Figures S5 and S6.

**Figure S5.**
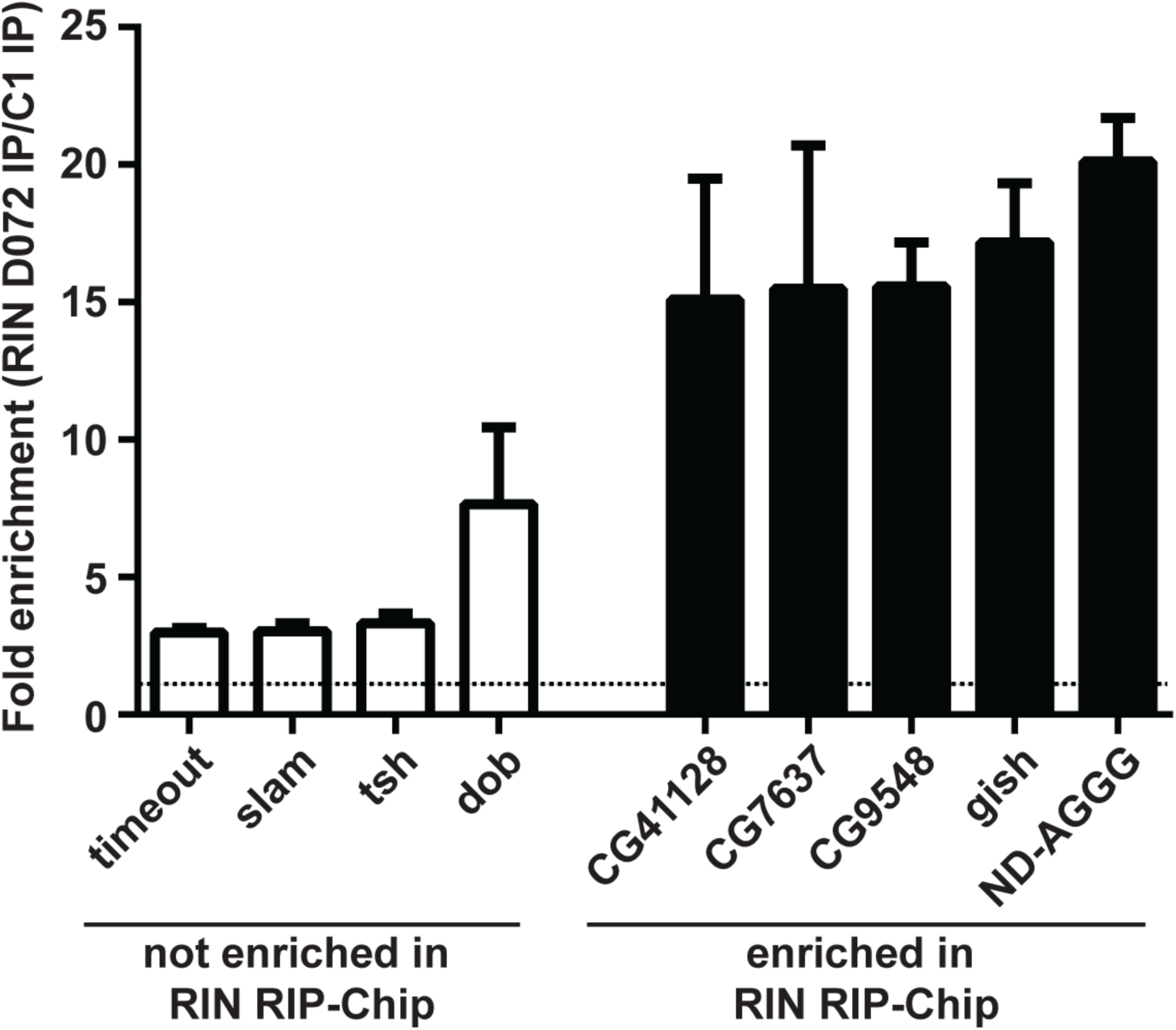
RT-qPCR validation of the enrichment of a subset of RIN-associated mRNAs identified by RIN (D072) RIP-Chip. Two biological replicates of RIN (D072) RIP and C1 RIP were performed, independent of the samples used for the RIP-Chip experiment, and RT-qPCR was used to quantify the levels of various mRNAs defined by RIP-Chip as RIN non-target mRNAs (*timeout*, *slam*, *tsh*, *dob*) or RIN-associated mRNAs (*CG41128*, *CG7637*, *CG9548*, *gish*, *ND-AGGG*). Shown are bar plots depicting the enrichment of mRNA levels in the RIN (D072) RIP vs C1 RIP. These enrichments observed by RT-qPCR are consistent with the RIP-Chip results. Error bars indicate standard deviation.

**Figure S6.**
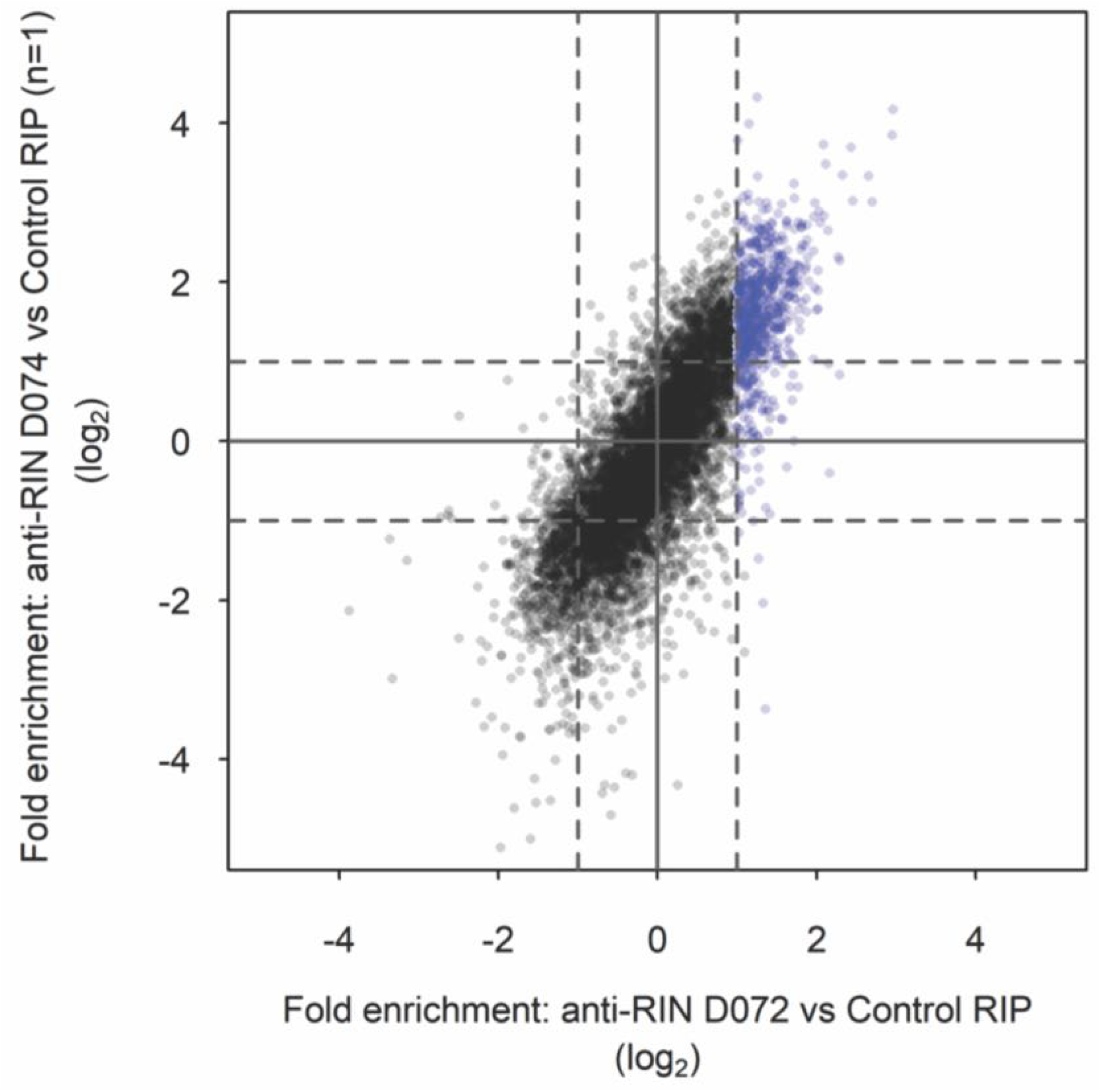
Correlation of RIN (D072) RIP-Chip fold-enrichment data with RIN (D074) RIP-Chip. To corroborate the RIN RIP-Chip data obtained using the anti-RIN D072 Fab, a single biological replicate of RIP-Chip was carried out using an additional anti-RIN Fab, D074, with the C1 Fab as a control. Shown are scatterplots depicting the relationship of fold-enrichments (level in RIN IP / level in C1 IP) between the RIP-Chip performed with the anti-RIN D072 Fab (average of three biological replicates) and the anti-RIN D074 Fab (single biological replicate), for all transcripts represented on the microarray that were defined as expressed in early embryos. There is a strong correlation between the fold enrichments obtained using the different anti-RIN Fabs. Transcripts defined in the anti-RIN D072 RIP-Chip as RIN-associated are highlighted in blue.

**Figure S7.**
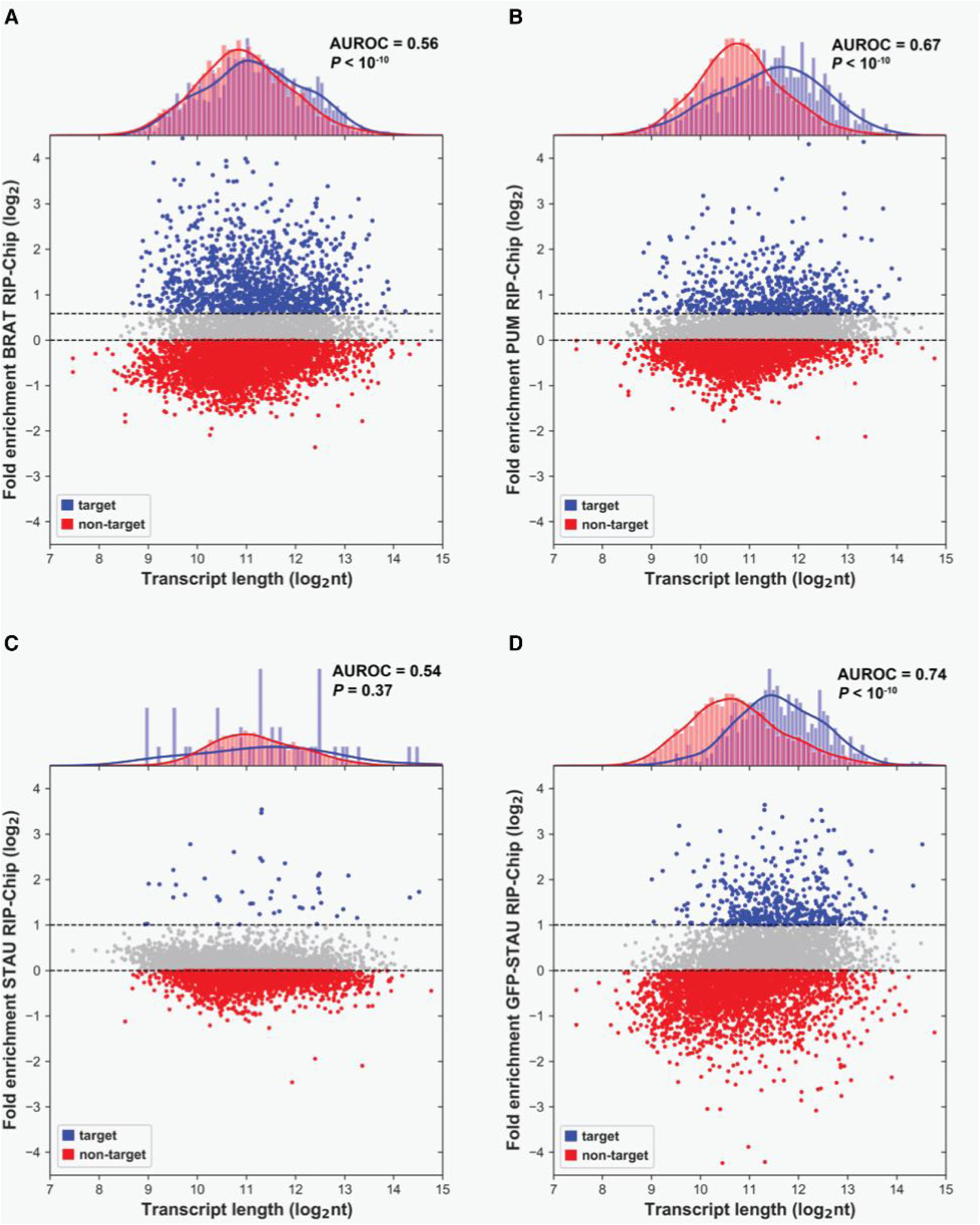
BRAT, PUM, and STAU-associated transcripts are not short. Plots showing the relationship between transcript length and association with **(A)** BRAT, **(B)** PUM and **(C, D)** STAU, for all genes with transcripts represented on the microarray that were defined as expressed in early embryos in previous studies (Laver et al., 2013 and 2015a). Transcript length for each gene was taken to be the length of the longest annotated transcript isoform, and RBP association is represented by the fold-enrichment in the RBP RIP-Chip versus control RIP-Chip of the most highly enriched probe set on the microarray for each gene. Genes encoding RBP-associated transcripts (defined as enriched > 1.5-fold for PUM and BRAT RIP and > 2-fold for STAU and GFP-STAU, versus control RIP with FDR < 5%) are highlighted in blue. Co-expressed non-targets (defined as log_2_ fold-enrichment in RBP versus C1 RIPs < 0 in all cases) are highlighted in red. Gene length distributions of RBP targets and non-targets are shown in the density plots at the top. AUROC and *P* value of Mann-Whitney *U* test on transcript lengths of RIN targets and non-targets is shown on each plot.

**Figure S8.**
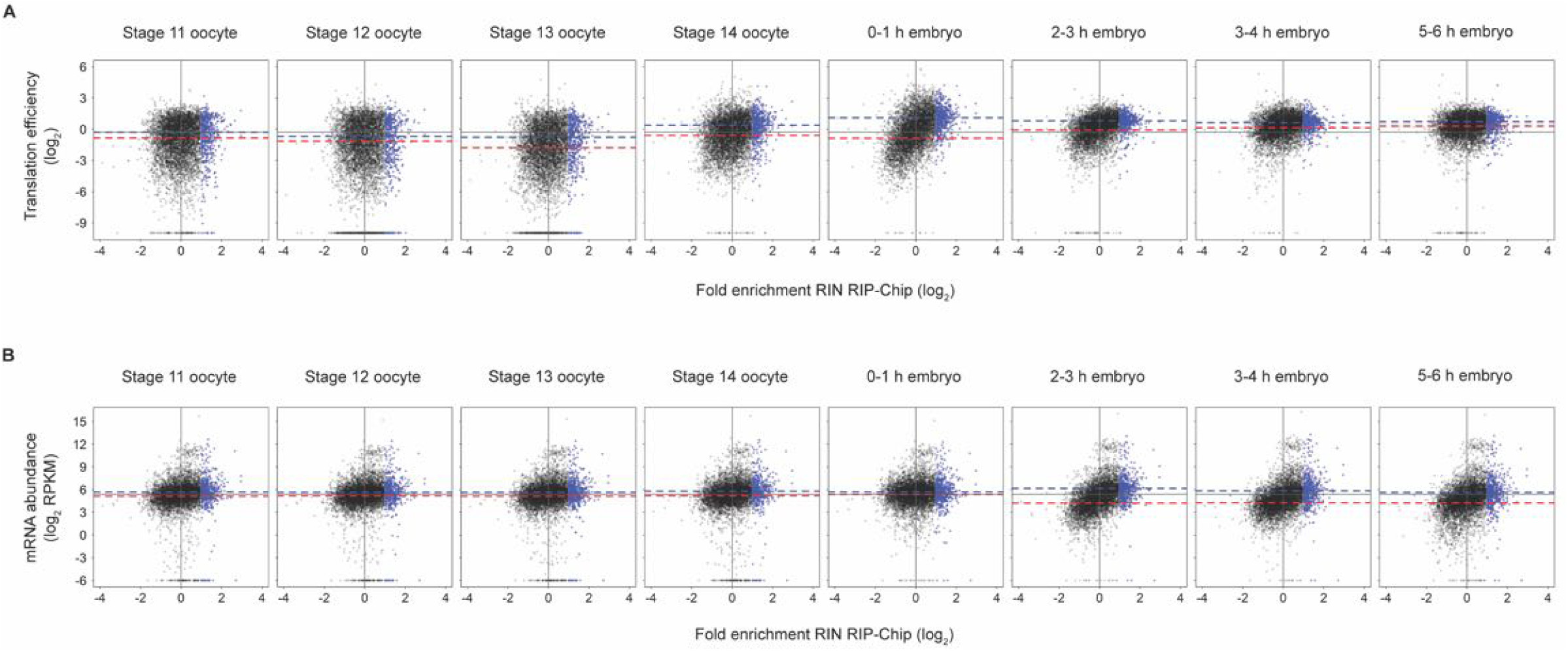
Scatterplots showing the relationship between association with RIN and translational efficiency or mRNA abundance throughout late oogenesis and early embryogenesis. Scatterplots showing the relationship between **(A)** translational efficiency, or **(B)** mRNA abundance, as reported by Eichhorn et al., 2016, and association with RIN. Points are shown for all genes that were measured in both this study and Eichhorn et al., 2016. RIN-associated transcripts as defined in this manuscript are highlighted as blue points, and solid vertical lines indicate no enrichment in RIN RIP (i.e. log_2_ fold enrichment = 0). Dashed blue horizontal lines indicate median values of translational efficiency (A), or mRNA levels (B) for RIN-associated transcripts at each timepoint. Dashed red horizontal lines indicate median values of translational efficiency (A), or mRNA levels (B) for non-RIN targets (defined as fold-enrichment in RIN/C1 RIPs < 1, or log_2_ of fold-enrichment < 0) at each timepoint. For (A) and (B), at all timepoints solid dark-grey horizontal lines indicate the median value of translational efficiency (A) or mRNA levels (B) in stage 14 oocytes as a reference.

**Figure S9.**
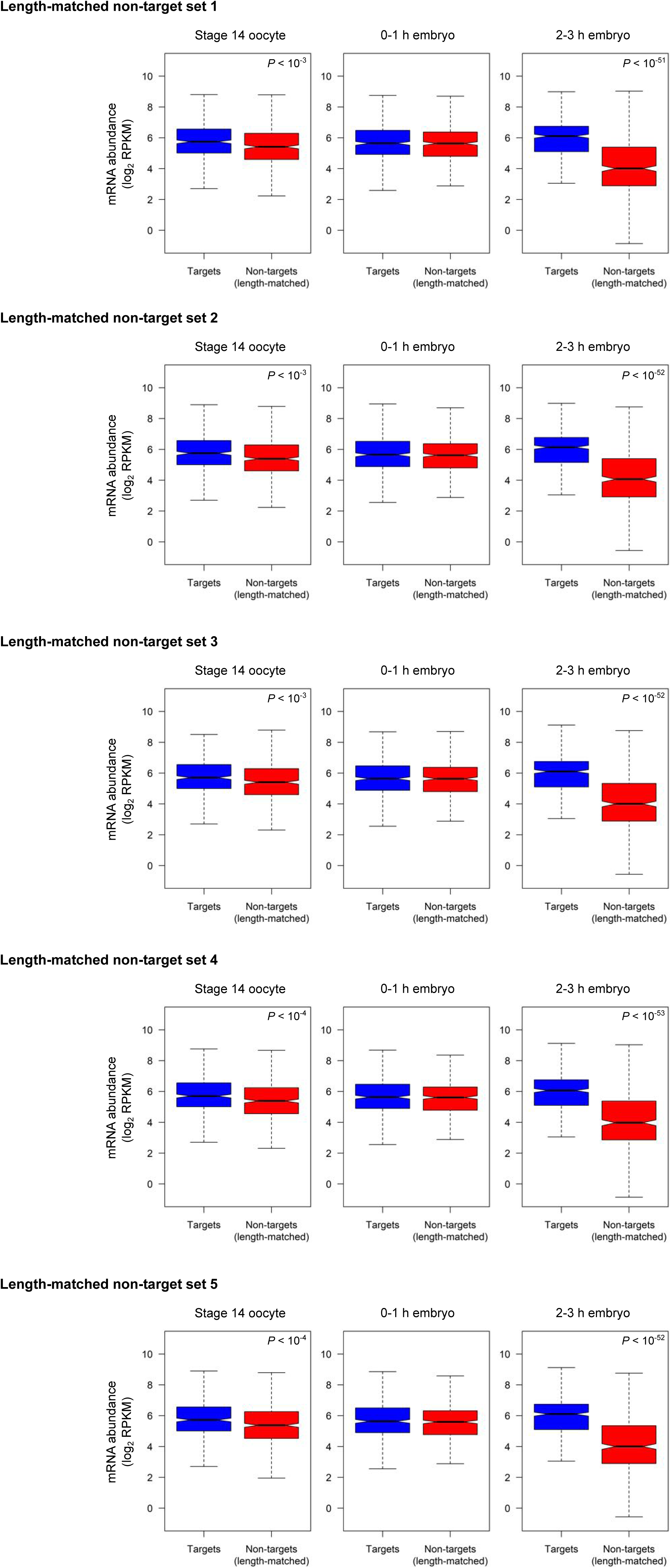
Boxplots comparing mRNA levels of RIN-associated mRNAs to length-matched non-RIN-targets. mRNA levels, measured by Eichhorn et al. (2016), of RIN-associated transcripts (“Targets”; blue) and a five sets of randomly selected, length-matched, co-expressed non-RIN-targets (red), for each time point. Data for the first set is also shown in Figure 3B. For comparisons with a statistically significant difference in mRNA levels between targets and length-matched non-targets, Wilcoxon rank-sum *P* values are indicated in the top right of the plots.

**Figure S10.**
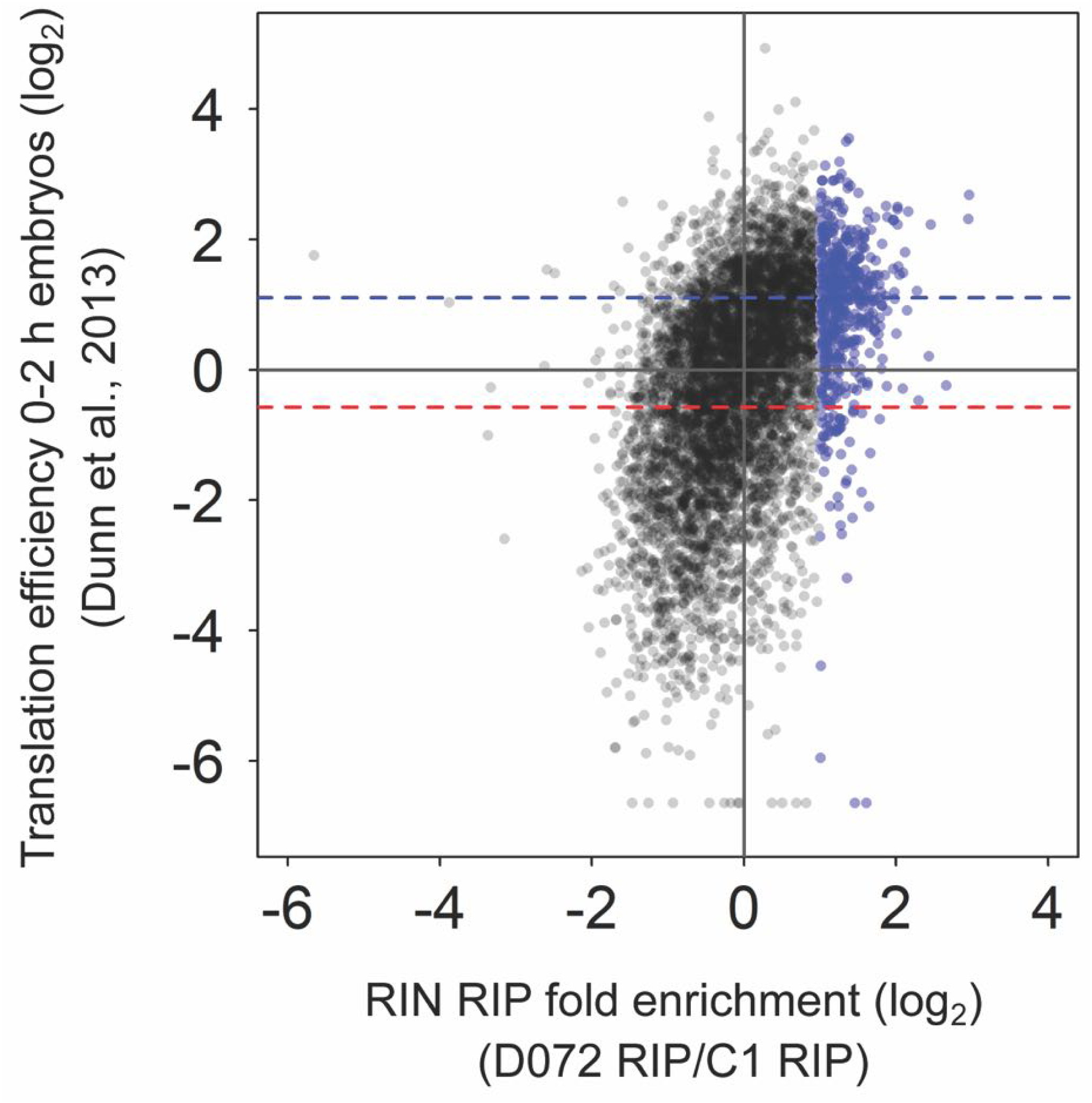
RIN-associated transcripts have higher levels of translation than RIN non-target transcripts in 0-2 h embryos, based on genome-wide translation efficiencies reported by Dunn et al., 2013. Scatterplot showing relationship between fold-enrichment in RIN (D072) RIP-Chip, and translational efficiency in embryos collected 0-2 hours post-egg-laying as reported in Dunn et al., 2013. Points are shown for all genes that were measured in both this study and Dunn et al., 2013. RIN-associated transcripts as defined in this study are highlighted as blue points. Dashed blue horizontal line indicates median translational efficiency for RIN-associated transcripts, and dashed red horizontal line indicates median translational efficiency for non-RIN targets (defined as fold-enrichment in RIN/C1 RIPs < 1, or log_2_ of fold-enrichment < 0). There is a significant correlation between enrichment in the RIN RIP and translational efficiency (Spearman rho = 0.47, *P* = 1.22 x 10^-310^), and RIN-associated transcripts have significantly higher translation efficiencies than RIN non-targets (Wilcoxon rank-sum *P* = 1.86 x 10^-113^).

**Figure S11.**
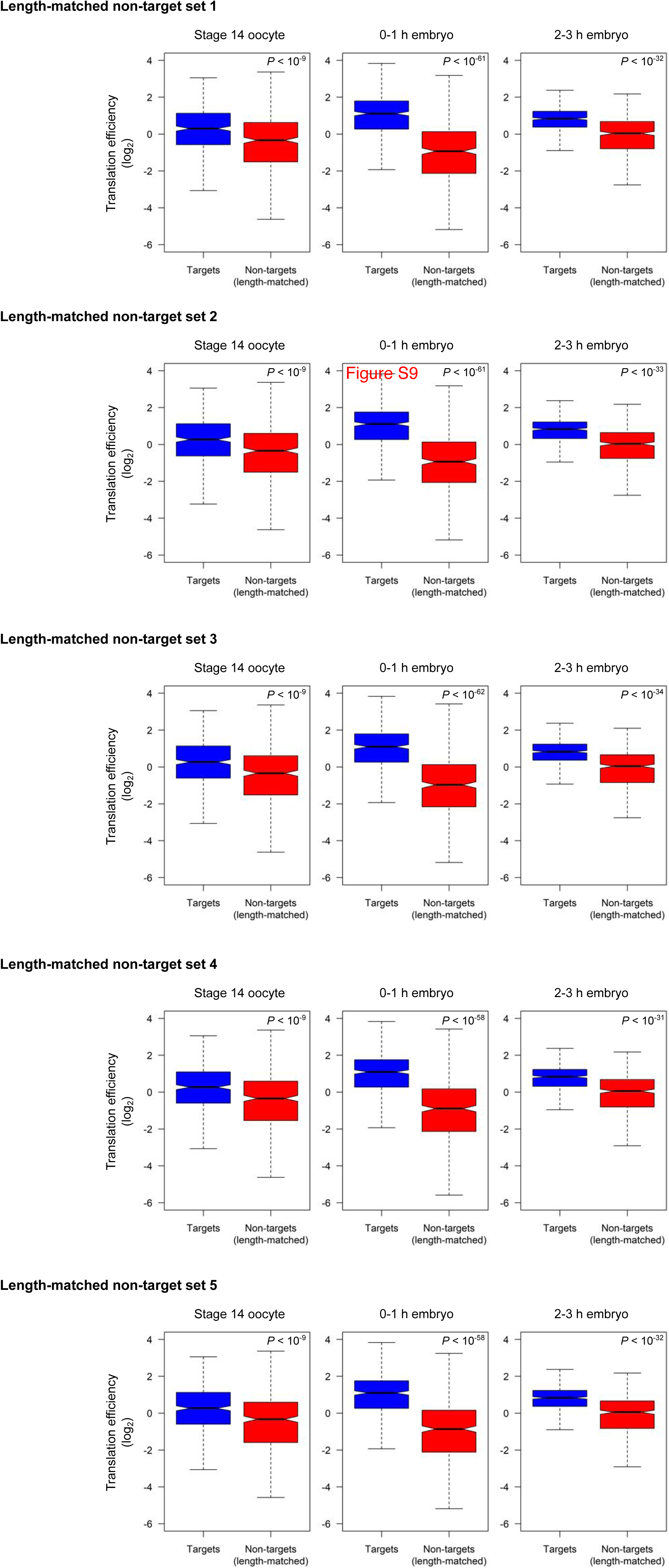
Boxplots comparing translational efficiency of RIN-associated mRNAs to length-matched non-RIN-targets. Translational efficiency, measured by Eichhorn et al. (2016), of RIN-associated transcripts (“Targets”; blue) and a five sets of randomly selected, length-matched, co-expressed non-RIN-targets (red), for each time point. Data for the first set is also shown in Figure 4B. For comparisons with a statistically significant difference in mRNA levels between targets and length-matched non-targets, Wilcoxon rank-sum *P* values are indicated in the top right of the plots.

**Figure S12.**
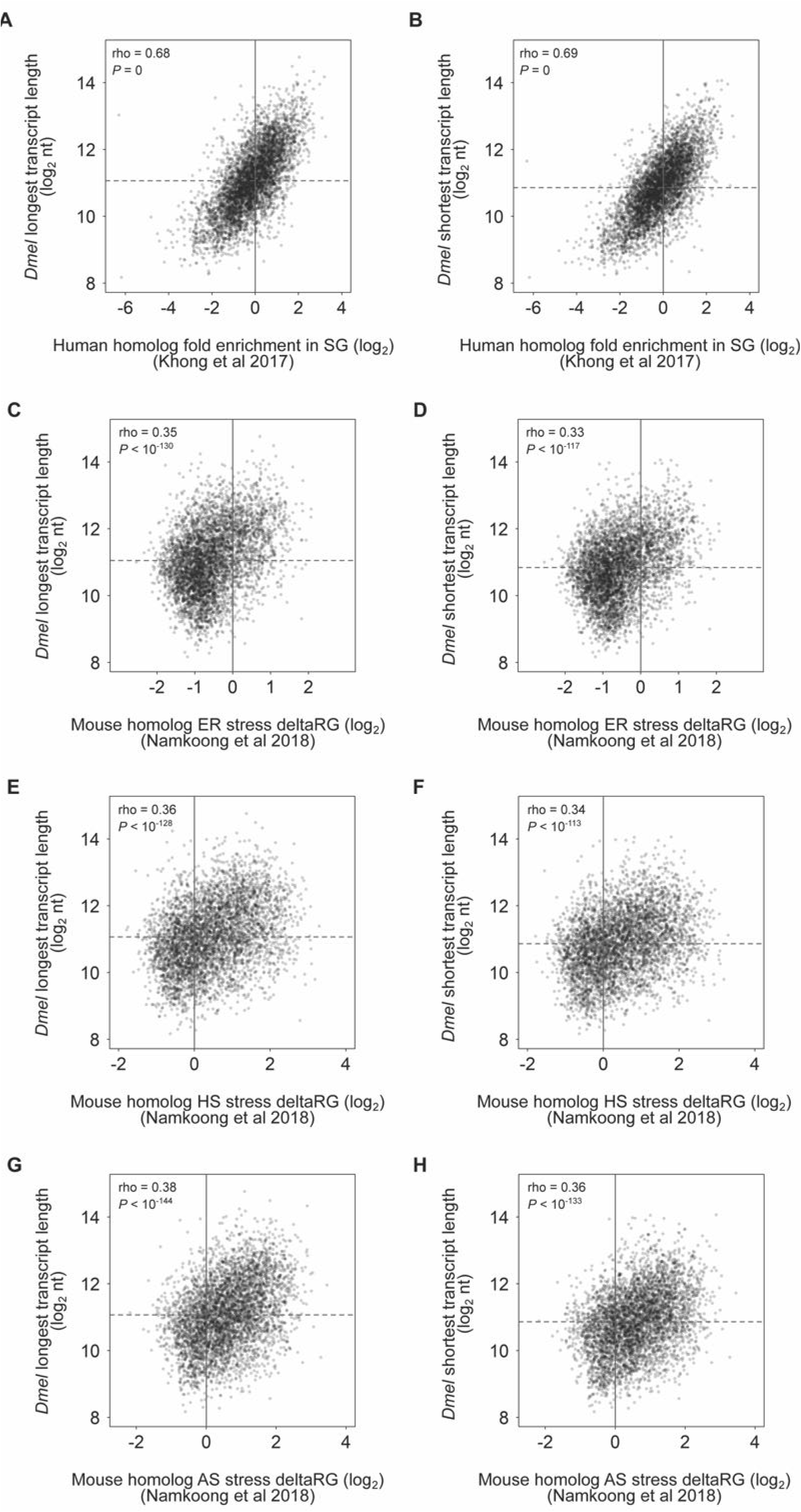
The fly homologs of SG-enriched or -depleted transcripts show the same length distribution as the mammalian transcripts. The transcript lengths of *Drosophila* genes were compared to published measurements of the degree to which their mammalian homologues are enriched in stress-granules, as defined for human homologues in Khong et al. (2017) **(A, B)**, or mouse homologues in Namkoong et al., 2018 **(C-H)**. In Khong et al., stress granule enrichment was measured as the amount of a given transcript present in isolated stress granule “cores” relative to total RNA, in response to arsenite stress. In Namkoong et al., stress granule enrichment was measured as the change in the percentage of a given transcript in the insoluble RNP granule fraction (“deltaRG”) in response to three different stresses: endoplasmic reticulum (ER) stress (panels C and D), heat shock stress (“HS”; panels E and F), and arsenite stress (“AS”; panels G and H). Transcript lengths of *Drosophila* genes were represented by the length of either the longest annotated transcript isoform (A, C, E, G) or the shortest annotated transcript isoform (B, D, F, H). Consistent with the relationship between transcript length and stress granule enrichment reported in Khong et al. and Namkoong et al., in all comparisons *Drosophila* genes with longer transcripts were more likely to have homologues enriched in stress granules, as indicated by the Spearman correlation rho values and associated *P*-values indicated on the plots.

**Figure S13.**
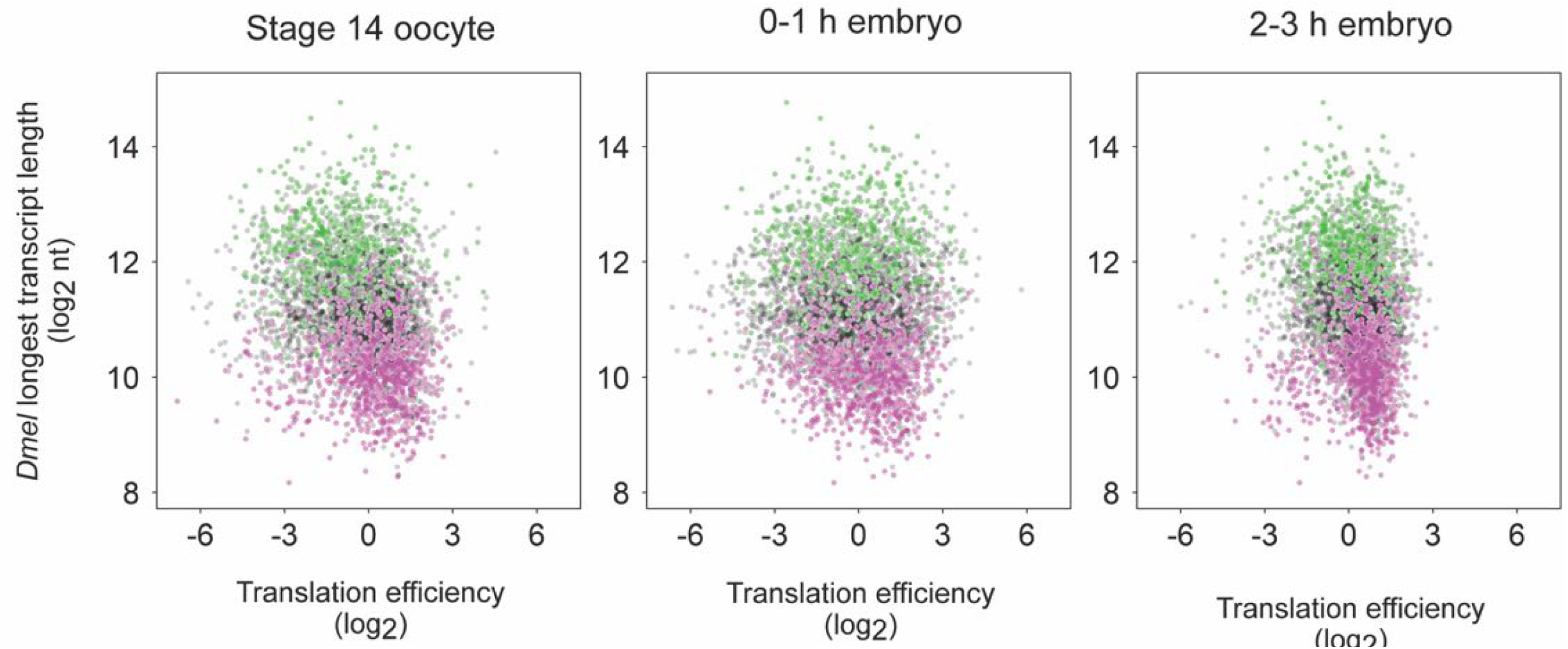
Relationship of transcript length and translational efficiency of *Drosophila* transcripts with stress granule-association status of their homologous mammalian transcripts. Plots of *Drosophila* transcript length (represented by length of the longest annotated isoform) versus translational efficiency (as measured in Eichhorn et al., 2016) are shown for translational efficiency values measured by Eichhorn et al. in Stage 14 oocytes, 0–1 h embryos, and 2–3 h embryos. Transcripts whose human homologues have been defined as enriched in stress granules, by Khong et al. (2017), are highlighted as green points, and transcripts whose human homologues have been defined as depleted in stress granules by Khong et al. are highlighted as magenta points. The plots only show transcripts that were considered in all three studies (i.e., defined in our study as expressed in early embryos, measured in Eichhorn et al., and with human homologues measured in Khong et al.).

**Figure S14.**
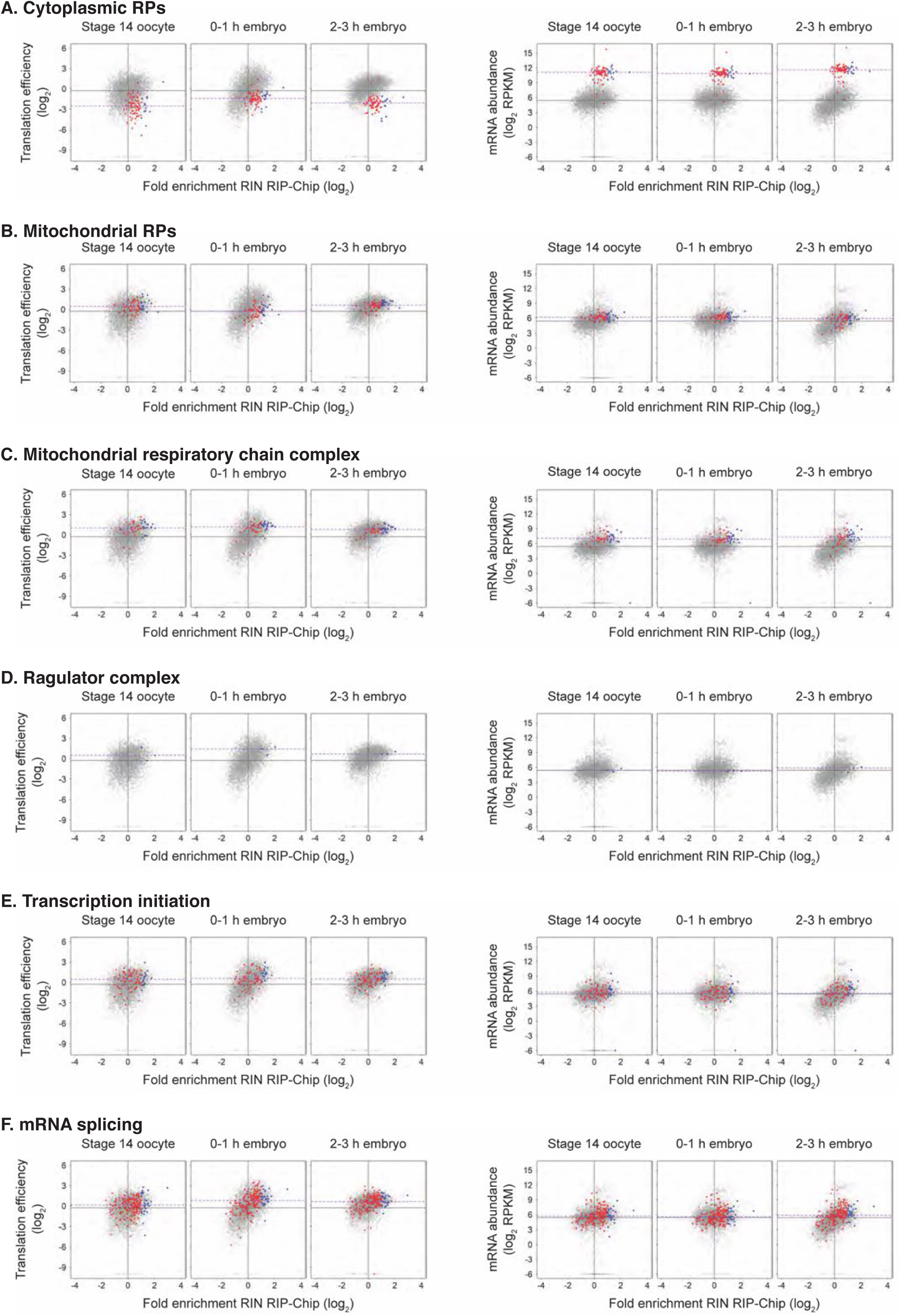
Relationship between RIN association, translational efficiency, transcript levels, and GO term annotations. Plots showing the relationship between fold-enrichment in the RIN RIP-Chip and translational efficiency (left-hand panels) or mRNA levels (right-hand panels), measured by Eichhorn et al. (2016), during timepoints spanning the maternal-to-zygotic transition, as plotted in Figure 3. Blue and red points represent genes annotated with the following GO terms, which are enriched among RIN-associated mRNAs: **(A)** cytoplasmic ribosomal proteins, **(B)** mitochondrial ribosomal proteins, **(C)** mitochondrial respiratory chain complex, **(D)** Ragulator complex, **(E)** transcription initiation, and **(F)** mRNA splicing. mRNAs we define as associated with RIN are highlighted in blue, and genes that fall below the cut-off used to define RIN-associated mRNAs are highlighted in red. Dashed purple horizontal lines indicate the median translational efficiency (left-hand panels) or mRNA levels (right-hand panels) for all genes annotated with a given GO term at each timepoint. For all timepoints, solid dark-grey horizontal lines indicate the median values of translational efficiency (left-hand panels) or mRNA levels (right-hand panels) in stage 14 oocytes, as a reference, and solid vertical lines indicate no enrichment in the RIN RIP-Chip. See Tables 1 and S5.

**Figure S15.**
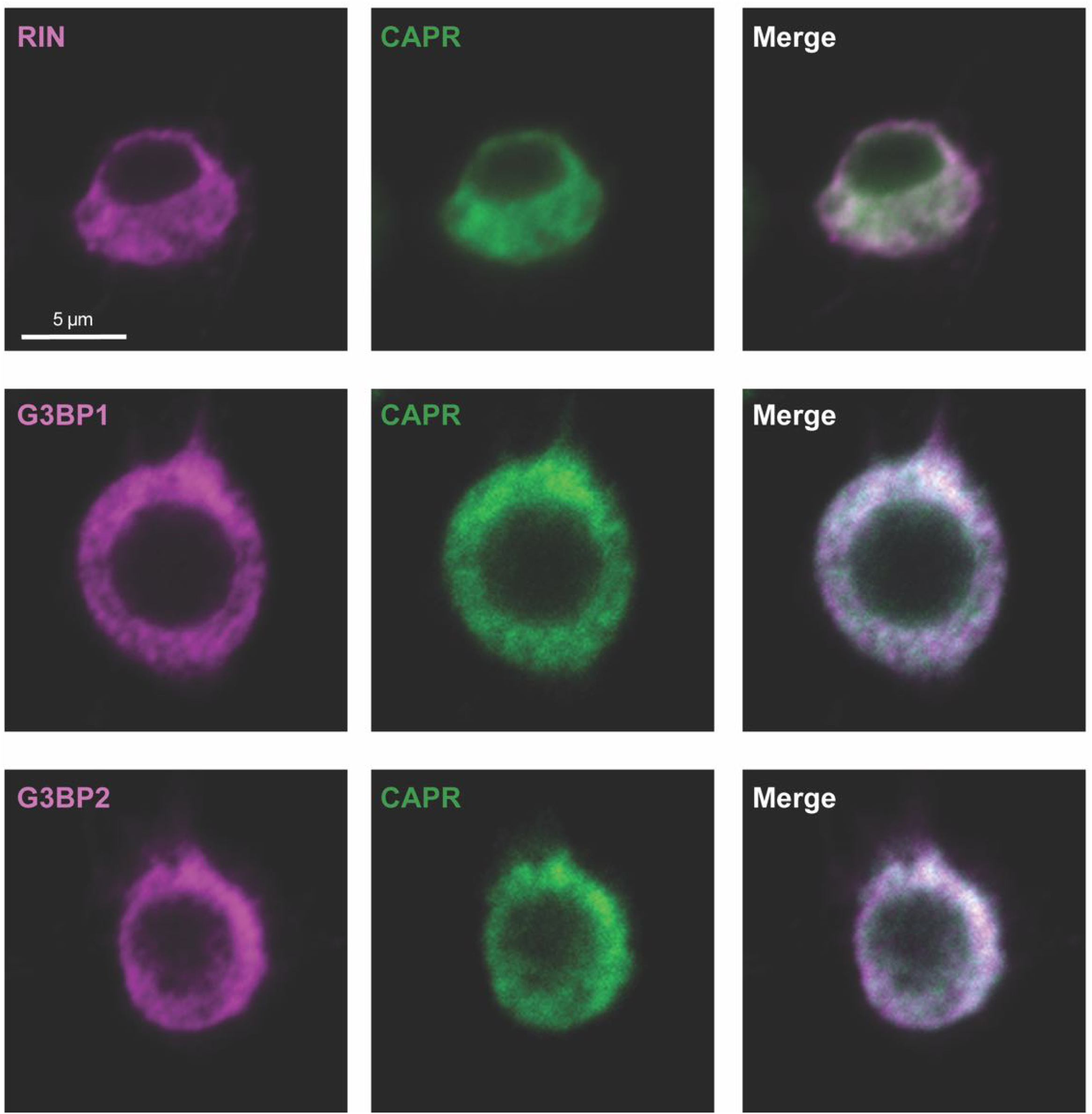
Distributions of RIN and CAPR in S2 tissue culture cells. Immunofluorescence visualization of **(A)** BIV-Tat-FLAG-RIN **(B)** BIV-Tat-FLAG-G3BP1, or **(C)** BIV-Tat-FLAG-G3BP2 tethered onto the Firefly luciferase 6x TAR reporter using anti-FLAG M2 in S2 cells. Co-staining with anti-CAPR (Papoulas et al., 2010) revealed that under conditions used for the tethering assay, RIN/G3BPs are ubiquitously distributed throughout the cytoplasm and partially colocalize with CAPR. In each of (A)-(C) at least 20 transfected cells were imaged and representative z-axis projections of three sections images are shown.

## Supplemental Tables

**Table S1. Results of IP-MS SAINT analysis for IPs (+)RNase and (–)RNase.**

(Excel spreadsheet)

**Table S2. Comparison of RIN interactors with published datasets on human and yeast G3BP/SGs**

(Excel spreadsheet)

**Table S3. List of RIN2-associated mRNAs with fold-enrichment and q-values.**

(Excel spreadsheet)

**Table S4. Comparison of transcript length for RBP target transcripts versus co-expressed non-target transcripts defined by RIP-Chip in previous studies**

(Excel spreadsheet)

**Table S5. Results of #ATS de novo motif discovery**

Table showing *de novo* motif discovery results with the discriminative motif finder #ATS (Li et al., 2010), which uses predicted sequence accessibility scores from RNAplfold (Bernhart et al., 2006) and identifies accessible sequence-specific RNA-binding protein motifs from *in vivo* binding data. Five samples comprised of the same number of length-paired targets and non-targets were extracted for each of the two different enrichment cut-offs (>2-fold and >3-fold) and regions (CDS and full transcript). Some targets were discarded during sampling because we could not find length-paired non-targets for them. We then performed *de novo* motif discovery with five-fold cross validation on each sample, but we failed to discover a consistent and significant motif in RIN targets.

(Excel spreadsheet)

**Table S6. Motif enrichment analysis**

(Excel spreadsheet)

**Table S7. Wilcoxon rank sum test *P* values for comparison of SRE scores of RIN targets vs non-targets**

(Excel spreadsheet)

**Table S8. Output from DAVID functional annotation enrichment analysis tool for RIN-associated mRNAs.**

(Excel spreadsheet)

**Table S9.**
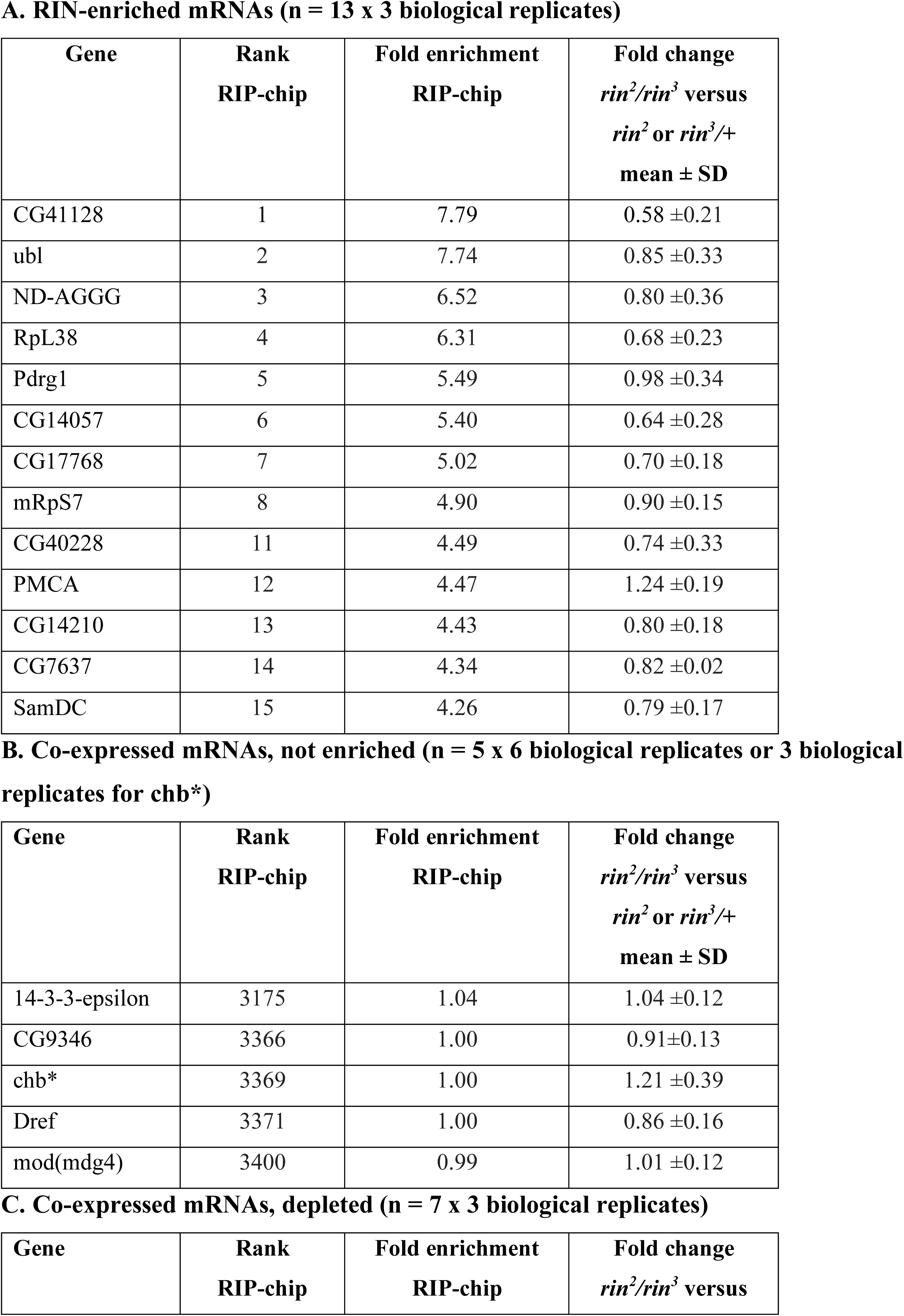
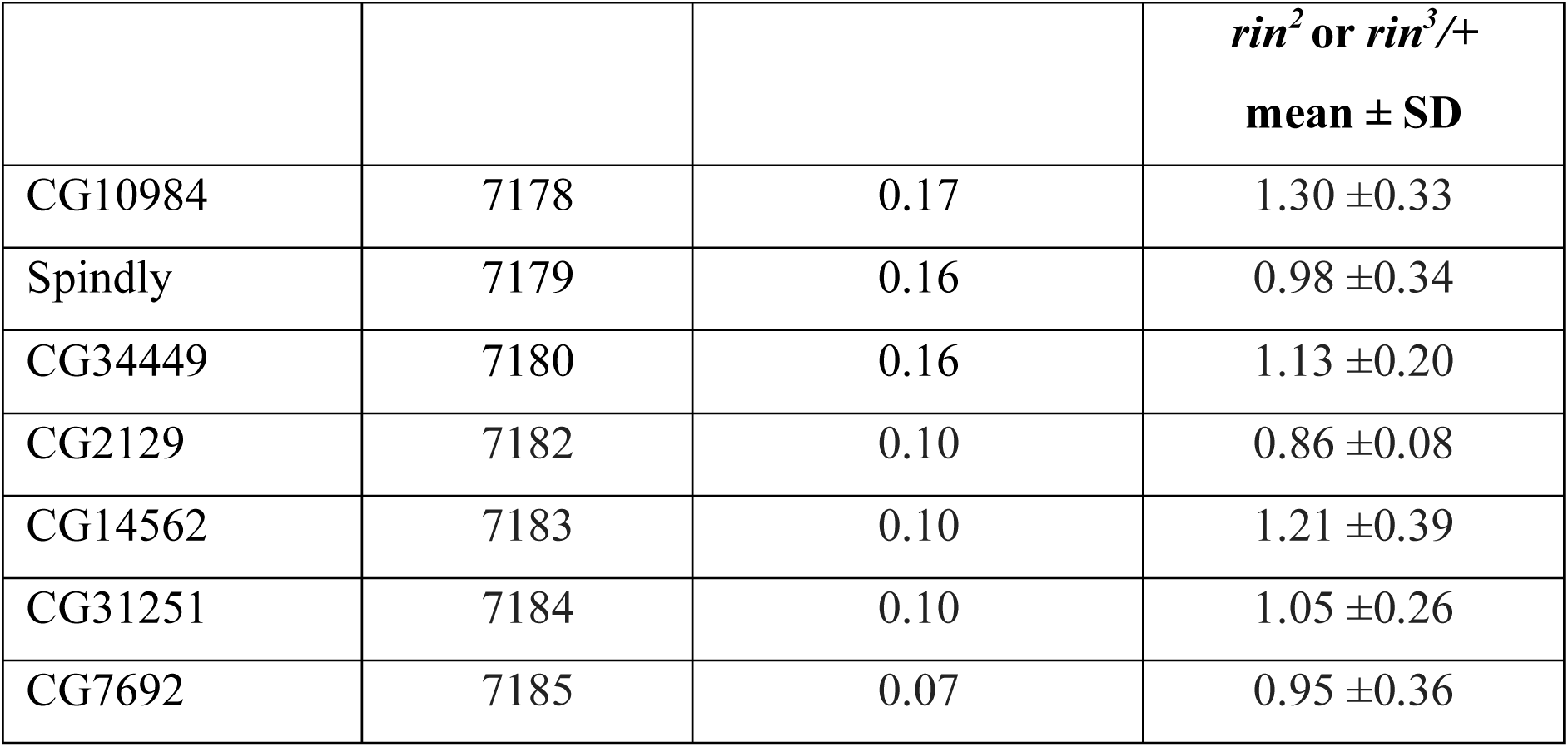
RT-qPCR of mRNAs from ovaries. A. RIN-enriched mRNAs (n = 13 x 3 biological replicates)

**Table S10.**
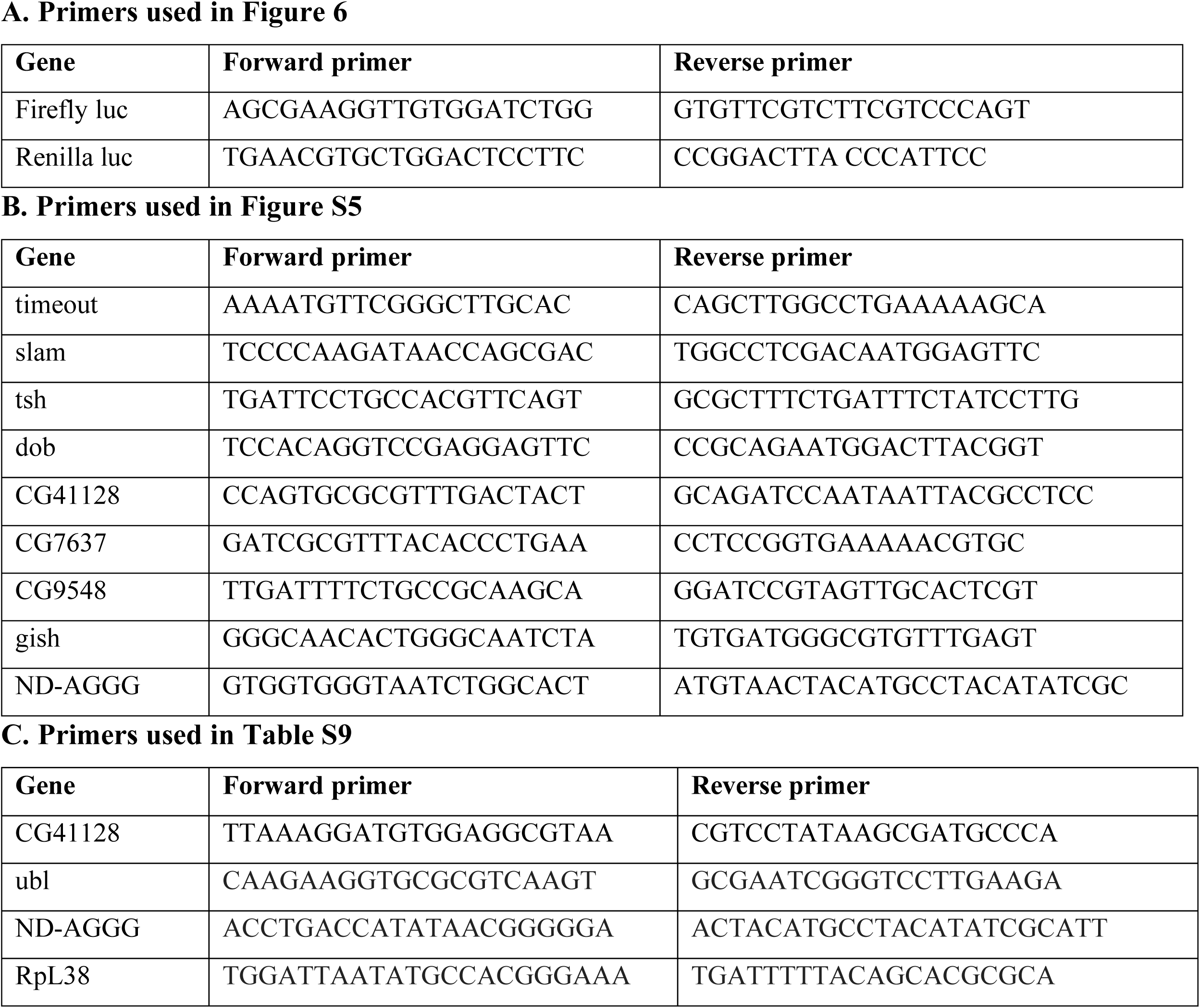
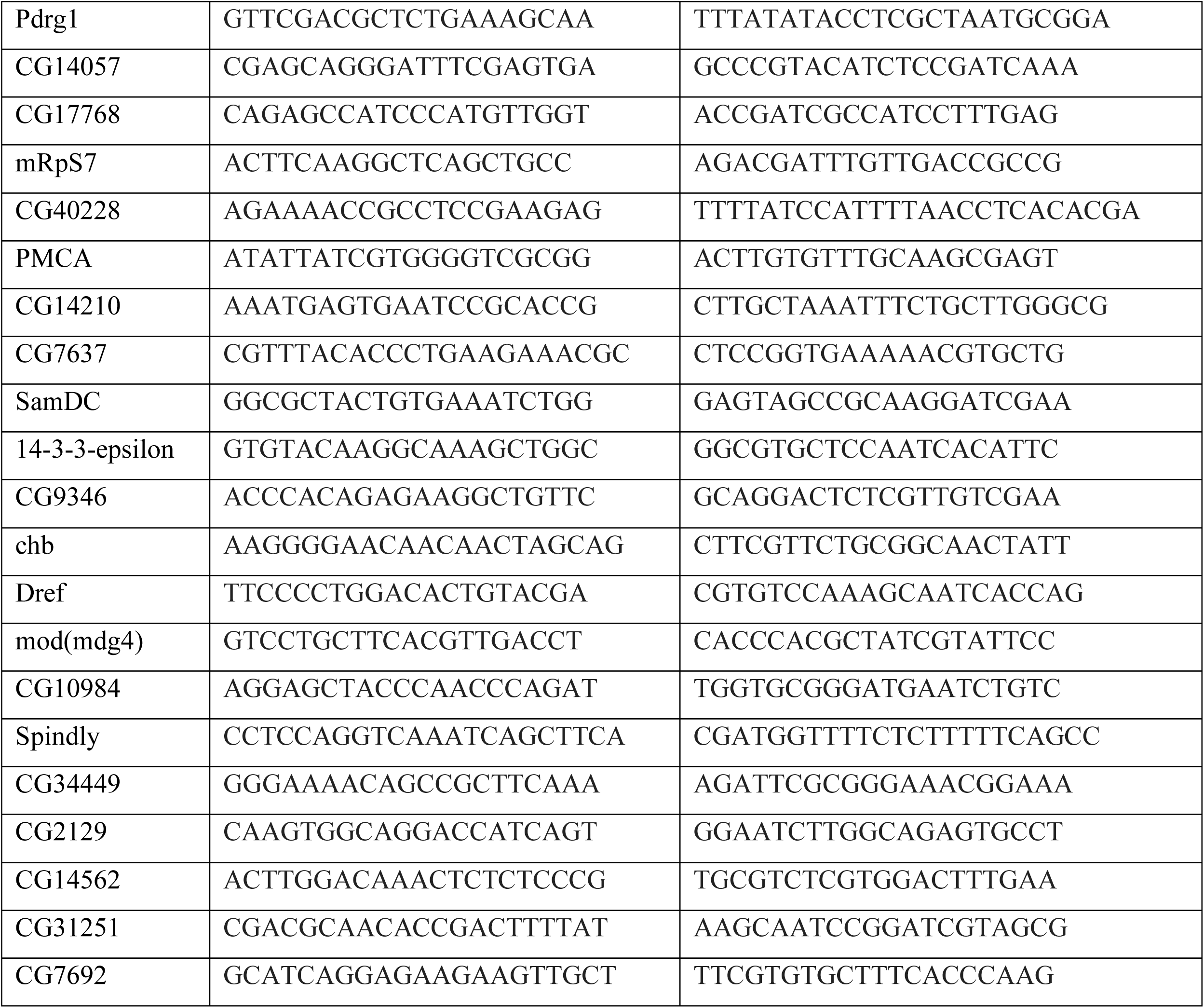
RT-qPCR primer sequences A. Primers used in Figure 6.

